# Tripartite host-parasite-virus interactions reshape chronic visceral leishmaniasis through persistent *Leptomonas seymouri* co-infection

**DOI:** 10.64898/2026.08.25.747179

**Authors:** Sayantan Das, Pritam Dey Sarkar, Rudra Chhajer, Subhajit Biswas

**Affiliations:** Infectious Diseases and Immunology Division, CSIR-Indian Institute of Chemical Biology, West Bengal, India; Academy of Scientific and Innovative Research (AcSIR), Ghaziabad-201002, India

**Keywords:** *Leishmania*, *Leptomonas*, macrophage, triple pathogen, Lepsey NLV1, kala-azar

## Abstract

**Background:** Visceral leishmaniasis (VL), caused by *Leishmania donovani* (LD), is increasingly associated with the insect-restricted trypanosomatid *Leptomonas seymouri* (LS), which harbours the RNA virus *Leptomonas seymouri* narna-like virus 1 (Lepsey NLV1). Our recent study demonstrated that LS co-infection with LD enhances survival of murine (RAW 264.7) and mammalian (THP-1) macrophages and augments LD and LS persistence compared to LD or LS mono-infection *in vitro.* However, the *in vivo* fate of LS and its viral endosymbiont during chronic VL remains poorly understood. This study investigated the long-term dynamics of parasite persistence, tissue dissemination and viral maintenance during experimental mono- and co-infection.

**Methods and Findings:** BALB/c mice were infected with LD, Lepsey NLV1-positive LS, virus-positive AG83 isolate, or LD: LS co-infections (2:1, 5:1 and 10:1) and monitored for up to seven months. Parasite burden, species composition and viral load were quantified using ITS1 qPCR, densitometry, nested RT-PCR and qRT-PCR, supported by microscopy and immunofluorescence assay. LS established productive visceral infection independently, with parasite burdens exceeding the infecting inoculum, indicating active *in vivo* replication. Co-infection, particularly at a 10:1 LD: LS ratio, promoted the greatest long-term parasite persistence in visceral organs. Temporal analysis revealed early predominance of LS followed by progressive recovery of LD during chronic infection. Lepsey NLV1 was detected in visceral organs and blood for at least up to five months. Morphological analyses demonstrated intracellular LS amastigote-like forms in murine macrophages and transformation of splenic parasites into promastigotes, confirming parasite viability within mammalian tissues.

**Conclusions:** These findings demonstrate sustained visceral persistence of Lepsey NLV1-positive LS in mice and identify dynamic host-parasite-virus interactions that reshape infection during chronic co-infection. This work challenges the conventional view of VL as a strictly mono-parasitic disease and highlights a previously underappreciated tripartite interaction with potential implications of LS and its virus endosymbiont for VL pathogenesis.

## Introduction

Visceral leishmaniasis (VL), caused predominantly by *Leishmania donovani* (LD), remains a major global heath burden, particularly in endemic regions of the Indian subcontinent (1,2). The disease is characterised by chronic infection of reticuloendothelial organs such as the spleen, liver, and bone marrow, where parasites establish long-term persistence despite active host-immune responses (3). While the pathogenesis of VL has been extensively studied in the context of LD mono-infections (4), accumulating evidence suggests that natural infections may involve complex parasite communities and associated endosymbionts, the biological consequences of which remain poorly understood (5).

*Leptomonas seymouri* (LS), a traditionally insect-restricted monoxenous trypanosomatid, has emerged as a surprising co-infecting organism in clinical and experimental settings (6,7). Another unresolved question concerns the capacity of monoxenous trypanosomatid to adapt to mammalian hosts (8). Although considered non-pathogenic to mammals, its repeated detection alongside LD in VL and PKDL cases challenges this assumption (9–11). Our recent findings had demonstrated that LS is not merely a passive participant but can actively infect, replicate and persist within mammalian macrophages, particularly in co-infection settings (12). Furthermore, the temporal dynamics of parasite competition, tissue tropism and systemic dissemination in mixed infections had not been systematically characterized *in vivo*.

Notably, LS, either alone or isolated from LD/PKDL clinical laboratory samples or serum from affected individuals, was shown to harbour an RNA virus, *Leptomonas seymouri* narna-like virus 1 (Lepsey NLV1) (13,14). This finding drew parallels with other trypanosomatid-associated viruses that were reported to modulate host immunity and parasite virulence (15,16). The contribution of viral endosymbionts to parasite fitness and host-pathogen interactions has gained increasing attention, particularly following studies on Leishmania RNA viruses (LRVs) in cutaneous and mucocutaneous leishmaniasis (17–20). Presence of virus within parasites can exacerbate disease severity by triggering host innate immune pathways, altering cytokine responses, and promoting parasite survival (21–23). Notably, our recent findings indicated that Lepsey NLV1 was persistently detectable within infected macrophages and exhibited significantly higher replication within LS, in co-infection scenarios compared to mono-infections. This suggests that parasite-virus interactions are enhanced in the presence of LD (12). More importantly, the tripartite interaction between a dixenous parasite (LD), a monoxenous parasite (LS), and its viral endosymbiont (Lepsey NLV1) represents an under-investigated paradigm with potential implications for disease heterogeneity and treatment outcomes.

In this study, we employed a murine model to investigate the long-term *in vivo* dynamics of co-infection between LD and LS harbouring Lepsey NLV1, focusing on parasite burden, viral persistence and host-parasite interactions across visceral organs and circulation. By integrating quantitative molecular approaches with microscopic and immunological analysis, we uncover distinct temporal shift in parasite dominance, sustained viral persistence and evidence for intracellular survival of LS within mammalian host tissues. Our findings provide novel insights into the role of viral endosymbionts in shaping co-infection biology. Collectively, this work advances our understanding of multi-pathogenic and virus-associated parasitic infections, highlighting a previously underappreciated layer of complexity in VL pathogenesis and opening new avenues for investigating host-parasite-virus interactions in chronic infectious diseases.

## Materials and methods

### 2.1. Parasite Culture

LD (MHOM/IN/1983/AG83), LS (ATCC) and a naturally mixed virus-positive AG83 laboratory isolate promastigotes were maintained at 22 °C in M199 medium (Sigma) supplemented with 10% heat-inactivated fetal bovine serum (FBS; Gibco) and 1% L-glutamine-penicillin-streptomycin solution (Sigma). Stationary-phase promastigotes were harvested on day 7 of culture and used for subsequent *in vitro* and *in vivo* infection experiments.

### 2.2. Experimental Animals

Male BALB/c mice (n = 40), aged 4-6 weeks and weighing 20-30 g, were housed in individually ventilated cages under controlled environmental conditions (23 ± 3°C; 55 ± 5% relative humidity) with a 12-hours (h) light/dark cycle. Animals had ad libitum access to standard laboratory chow diet and drinking water throughout the study. Mice were acclimatized to the experimental facility for at least 7 days prior to study initiation. All the animals were acclimatized for at least 7 days before the initiation of the experiment. The animals were maintained under standard housing conditions throughout the study and euthanized at the end of the experiment according to the study protocol for further analysis.

### 2.3. Ethics statement

All experimental protocols were performed in accordance with the Committee for the Control and Supervision of Experiments on Animals (CCSEA) guidelines. The study protocol was designed based on literature review in the same area and it was approved by Institutional Animal Ethics Committee (IAEC, CSIR-IICB, Kolkata, Approval identifier: IICB/AEC/Meeting/Sep/2023/6) (24,25). The experiments on animals comply to the ARRIVE guidelines.

### 2.4. *In vivo* Infection

Balb/c mice (n=3-5/ group) were infected with 2 × 10^7^ stationary phase parasites [LD, LS, virus positive AG83 lab isolate and 2:1, 5:1 or 10:1 co-infection (LD: LS)]. Un-infected mice were maintained as experimental control. The parasite inoculum was selected based on previously published studies (24,25). Prior to infection, ITS1 PCR analysis of the virus-positive AG83 laboratory isolate revealed an LD:LS ratio of approximately 7:1. Therefore, a 10:1 (LD:LS) ratio was included in majority of co-infection experiments to represent a comparable parasite ratio to that observed in the laboratory isolate. 100 uL parasite inoculum was injected intravenously (I.V.). Only virus positive AG83 lab isolate was injected via intraperitoneal (I.P.) route. Infection was carried out for different time points ranging from 1-7 months post infection (m.p.i). On sacrifice, spleen, liver and blood samples were collected for further analysis.

### 2.5. DNA extraction and ITS-1 gene PCR

For *in vivo* analysis, DNA was extracted from mice blood, liver, spleen tissue and splenic tissue cultures by means of DNeasy Blood and Tissue Kit (Qiagen), following manufacturer’s protocol. Approximately, 10 mg of spleen and 20 mg of liver tissue were vortexed and triturated in warm tissue lysis buffer (provided in the kit) until the tissue was completely lysed before processing for down-stream DNA extraction. For splenic tissue cultures, 500 uL cultures were subjected to DNA extraction. The quality and concentration of DNA was assessed using a NanoDrop spectrophotometer (Thermo Scientific, USA).

LS DNA specific PCR product or rDNA-ITS1 fragment was directly amplified from the isolated DNA using respective forward and reverse primers and protocol as described previously (12,14).

### 2.6. Quantitative Real-Time PCR and Densitometric Analysis

Parasite burden was quantified using a SYBR Green-based quantitative real-time PCR (qPCR) assay performed with the Luna Universal PCR Master Mix (New England Biolabs). Equal amounts of genomic DNA from each experimental condition were used as input to ensure comparability across samples. Amplification targeted the internal transcribed spacer 1 (ITS1) region, generating species-specific fragments of 320 bp or 418 bp using previously validated primer sets (12,26). Quantitative amplification was carried out on a StepOne™ Real-Time PCR System (Applied Biosystems, Thermo Fisher Scientific) under optimized cycling conditions.

Amplification specificity was confirmed by resolving qPCR products on 1% agarose gels, verifying the presence of bands corresponding to the expected sizes of the PCR products. To estimate the relative abundance of LD and LS in co-infected samples, gel images were subjected to densitometric analysis using ImageJ software. Band intensities from experimental samples were quantified, enabling calculation of LD: LS ratios based on normalized signal intensities standard curve.

### 2.7. RNA Extraction, cDNA Synthesis, and Virus Detection

To assess virus burden *in vivo*, total RNA was extracted from blood, spleen, and liver tissues using the QIAamp® RNeasy Mini Kit (Qiagen), following the manufacturer’s protocol. RNA concentration and purity were evaluated using a NanoDrop spectrophotometer to ensure suitability for downstream applications.

Complementary DNA (cDNA) was synthesized from purified RNA, followed by virus-specific nested PCR amplification using previously described primers and protocols (15). Amplified products were resolved on 1% agarose gels to confirm the presence of expected PCR products. Bands corresponding to the target size were excised and purified using the QIAquick® PCR Purification Kit (Qiagen, Germany) prior to downstream sequencing analysis.

### 2.8. Quantitative Real-Time PCR for Virus Load Estimation

Virus RNA copy numbers in extracted nucleic acid samples were quantified using quantitative reverse transcription PCR (qRT-PCR). Amplification targeted a 338 bp region of the Lepsey NLV1 genome using the CT2-F and CT2-R primer set, following previously established protocols (15).

Amplified products were resolved on 1% agarose gels and visualized using SYBR Safe™ nucleic acid staining to confirm specificity and expected product size.

### 2.9. Sequence Analysis

Nucleotide sequences were obtained through bidirectional DNA sequencing of the purified PCR products, utilizing the same primers as in the initial PCR for sequence validation (15). Sequence accuracy was ensured by overlapping reads from at least one forward and one reverse primer, followed by alignment using MEGA X (27) and BioEdit software (version 7.2.5). The Lepsey NLV1 sequences were compared with other Lepsey NLV1 sequences available in NCBI GenBank [Accession numbers: KU935604.1 and KY628364.1].

### 2.10. Peritoneal macrophage isolation and *in vitro* infection

Primary peritoneal mouse macrophages (PMM) were obtained from mice by intraperitoneal administration of 1X ice-cold phosphate-buffered saline (PBS) with 1.4% Glucose. After isolating PMM, primary cells were centrifuged at 300g for 15min at 4 °C.

For *in vitro* infection assays, cells were resuspended in RPMI-1640 supplemented with 10% FBS and 1.5% L-Glut-pen-strep. 10^5^ cells were plated in a 4-well chamber (Genetix Biotech) tissue culture slide and incubated at 37 °C with 5% CO2 overnight. Infection was carried out according to previously standardized protocol (12) and incubated for 48-72 hours post infection (h.p.i.). After the incubation period, cells were fixed with methanol and stained with Giemsa solution (Himedia). Stained slides were examined under an optical microscope (EVOS) for determining the infection of macrophages at 1000x magnification.

### 2.11. Transformation of splenic parasites and Giemsa Staining

Spleens were aseptically harvested from infected BALB/c mice at 5 m.p.i. and processed under sterile conditions. Excised splenic tissues were gently minced and transferred to Schneider’s drosophila medium supplemented with 10% heat-inactivated fetal bovine serum (FBS) and 1% L-glutamine-penicillin-streptomycin. Tissue explants were maintained at 22 °C to facilitate parasite transformation and differentiation. Cultures were monitored periodically for parasite outgrowth.

After 14 days of *in vitro* culture, aliquots of the reactivated cultures were collected, and thin smears were prepared on clean glass slides. The smears were air-dried, fixed with methanol, and stained with Giemsa solution following standard protocols.

For histological stamp smears, small piece of spleen and liver from mice 7 m.p.i with LD, LS, virus-positive AG83 lab isolate, 10:1 Co-culture (LD: LS) and PBS control were dabbed on a glass slide and fixed with methanol followed by Giemsa staining. The stained slides were visualized under an optical microscope (EVOS) to assess parasite morphology.

### 2.12. Generation of mouse anti-*L. donovani* and anti-*L. seymouri* immune serum

Anti-*Leishmania donovani* (anti-LD) and anti-*Leishmania seymouri* (anti-LS) immune serums were generated in 6-week-old male BALB/c mice via I.V. administration of 2 × 10^7^ viable stationary-phase promastigotes of LD or LS. At 7 m.p.i., mice were euthanized, and blood samples were collected and centrifuged to obtain the corresponding antiserum.

### 2.13. Reactivity of mouse anti-LD and anti-LS immune serum

An indirect immunofluorescence (IF) assay was performed to evaluate the reactivity and cross-reactivity of mouse anti-LD and anti-LS immune serum. For murine macrophage (RAW 264.7) infection, 2 × 10 cells were cultured on 18 mm coverslips at 37 °C with 5% CO . Adherent macrophages were subsequently infected with stationary-phase LD or LS promastigotes at an MOI of 10 under the same incubation conditions. Uninfected RAW 264.7 cells were maintained as controls. After 4 h, non-internalized promastigotes were removed by washing with warm 1x PBS, followed by the addition of fresh DMEM-10. Infected as well as control cells were then incubated for 48 h.p.i. at 37 °C with 5% CO . To assess cross-reactivity, LD-infected macrophages were incubated with anti-LS serum, while LS-infected cells were incubated with anti-LD serum.

Briefly, infected macrophages containing intracellular amastigotes were washed twice with 1x PBS and fixed with 4% paraformaldehyde for 15 minutes at room temperature. Cells were then permeabilized using 0.1% Triton X-100 in PBS for 15 minutes. Following permeabilization, blocking was carried out for 1 h at room temperature using PBS containing 0.1% Triton X-100, 1% bovine serum albumin (BSA, sigma), and 22.5 mg/ mL glycine. Subsequently, cells were incubated with mouse anti-LD or anti-LS immune serum at a dilution of 1:10 for 1 h at room temperature. After washing, cells were treated with Alexa Fluor 488-conjugated anti-mouse IgG secondary antibody (ab150113) at a dilution of 1:500 for 1 h at room temperature in the dark, mounted by DAPI (ProLong TM Diamond Antifade DAPI; P36962). Finally, fluorescence visualization was performed using a Leica super-resolution confocal microscope.

### 2.14. Immunoblotting for cleaved caspase 3

A total of 3 × 10^5^ RAW 264.7 murine macrophage cells per well were seeded in a 12-well plate and were infected with LD, LS and co-culture (10:1). Infection was carried out according to previous protocol (12). At 72 h.p.i, cells were harvested in RIPA cell lysis buffer containing Protease inhibitor cocktail (Pierce Thermo fisher Scientific, A32955) and incubated for 10 min at 4 °C. It was followed by a centrifugation at 14000g for 10 min at 4 °C to remove the cell debris. The clear supernatant was collected and utilized for Western blot analysis. Protein concentration was determined using the BCA assay kit (Pierce).

Cell lysates were resolved on 5% stacking and 15% resolving SDS-PAGE gel using standard running buffer at 90V for 2.5 h. Protein ladders (Page Ruler, Thermo Scientific, 26616) were used for reference. Proteins were transferred to a nitrocellulose membrane via the Trans-Blot Turbo system (Bio-Rad) and blocked with 5% skimmed milk in TBST for 1 h at room temperature. To detect the target proteins, anti-cleaved caspase 3 antibody (CST-9661) was used at a dilution of 1:1000, β-Tubulin (CST-2128) was used as a housekeeping control diluted at 1:1000. Membrane strips containing cleaved Caspase 3 or GAPDH were incubated with the respective primary antibodies overnight at 4 °C in fresh TBST buffer. This was followed by incubation with secondary anti-rabbit IgG-HRP antibody (Abcam-97051) at 1:12000 dilution for 1 h at 4 °C. Protein bands were then visualized using ECL substrate (Bio-Rad) on the ChemiDoc system (Bio-Rad). Densitometric analysis was done using Image Lab software (Bio-Rad).

### 2.15. IL-12 (p40) ELISA

Quantitative IL-12 (p40) ELISA (BD OptEIA™, Cat. No.- 555165) was performed as per manufacturer’s protocol to determine cytokine levels from mice serum samples at 1,3 and 5 m.p.i. Standard curve was prepared using dilutions of the respective recombinant cytokine provided in the kit.

### 2.16. Statistical Analysis

All statistical analyses were conducted using GraphPad Prism version 8.0 (GraphPad Software, USA). Data are presented as mean ± standard deviation (SD). Appropriate statistical tests were selected based on the experimental design and data distribution. For comparisons involving more than two groups, one-way ANOVA followed by Dunnett tests were used. Each experiment included a minimum of three technical replicates. Statistical significance was defined as p < 0.05.

## RESULTS

### 1. Experimental murine infection reveals the *in vivo* infectivity of Lepsey NLV1-positive LS alone and in co-infection with LD

#### 1.1. Quantitative analysis reveals temporal remodeling of parasite burden and LD: LS dominance during chronic murine co-infection

Qualitative and quantitative ITS1 PCR analysis of visceral organs from mice infected with LD, LS or LD-LS co-cultures (2:1, 5:1, 10:1 or virus-positive AG83 lab isolate) confirmed the presence of parasitic DNA. At 3 m.p.i., LS mono-infected mice harboured parasite burdens exceeding the initial infecting inoculum (2 ×10^7^ parasites/ mouse) by approximately ∼3-fold in the spleen (6.8 × 10^7^ parasites/ spleen) and 265-fold in the liver (5.3 ×10^9^ parasites/ liver), demonstrating active *in vivo* replication and productive visceral infection independent of LD. Comparable or relatively higher parasite burdens were observed in both LD-LS co-infection groups (5: 1 and 10:1 respectively) compared to LS mono-infection, indicating that co-infection did not impair LS replication rather supported sustained parasite expansion in visceral tissues. LD mono-infected mice similarly exhibited robust tissue parasite burden (1.1 ×10^8^ parasites/ spleen and 4.1 × 10^9^ parasites/ liver), with all infection groups showing greater parasite accumulation in the liver than in the spleen **(Fig. 1A, 1B and Supplementary Table S1).**

**Figure 1.**
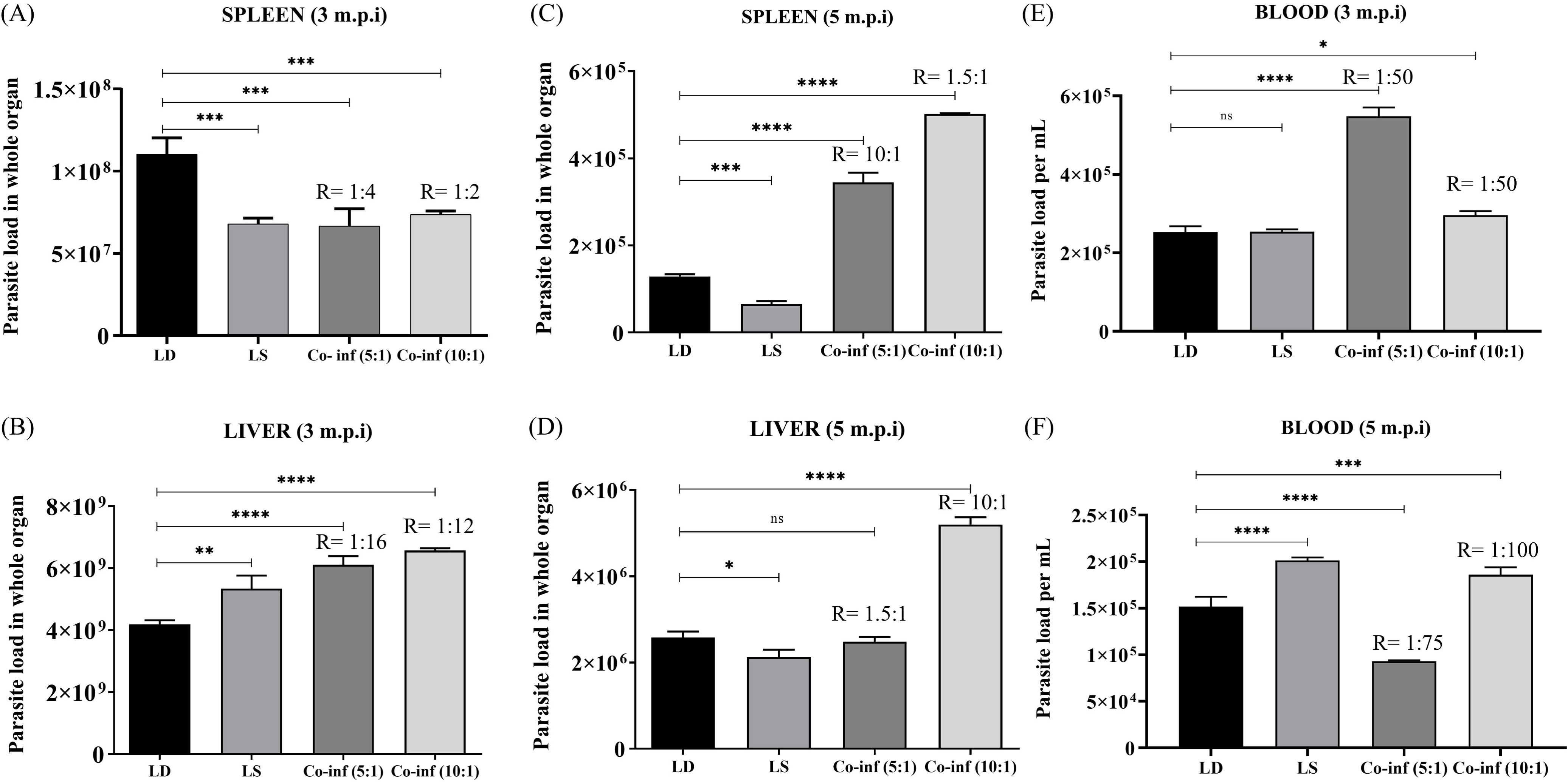
Temporal analysis of parasite burden in parasite infected BALB/c mice. Parasite burden was quantified by qPCR in BALB/c mice infected with LD, LS, or LD-LS co-infections at the indicated post-infection time points. **(A)** Spleen parasite burden at 3 months post-infection (m.p.i).; **(B)** Liver parasite burden at 3 m.p.i.; **(C)** Spleen parasite burden at 5 m.p.i.; **(D)** Liver parasite burden at 5 m.p.i.; **(E)** Blood parasite burden at 3 m.p.i.; and **(F)** Blood parasite burden at 5 m.p.i. The figure compares parasite persistence and temporal changes in parasite burden among the different infection groups across tissues. ‘R’ represents the LD: LS ratio derived upon ImageJ analysis. The data were represented as mean ± SD of a minimum of three independent experiments. Statistical significance was determined using one-way ANOVA followed by Dunnett test where significance is denoted as p < 0.05 (*), p < 0.01 (**), p < 0.001 (***), p < 0.0001 (****), ns = not significant.

These observations were further supported in an independent experiment using the virus-positive AG83 isolate. At 3 m.p.i., the liver parasite burden was estimated at (1.6 × 10^8^ parasites/ liver, LD: LS at 1: 10^4^), corresponding to a 7.8-fold increase relative to the infecting inoculum (2 × 10^7^ parasites/ mouse), whereas the spleen contained only (4.5 × 10^5^ parasites/ spleen, LD: LS at 1; 200), substantially below the inoculum. The increase in liver parasite burden beyond the infecting dose provides additional evidence of active *in vivo* replication (Supplementary Table S1).

By 5 m.p.i., LD, LS and both co-infection groups exhibited persistent infection. Although parasite burden declined markedly in both spleen and liver across all infection groups, LS remained readily detectable in both organs, confirming its long-term persistence in murine visceral organs. Despite the overall reduction in tissue parasite burden, the 10:1 LD: LS co-infection group retained the highest parasite burden in both spleen (5.0 × 10^5^ parasites/ spleen) and liver (5.1 × 10^6^ parasites/ liver), suggesting enhanced parasite persistence during co-infection. In contrast, LD (1.3 × 10^5^ parasites/ spleen and 2.5 × 10^6^ parasites/ liver) and LS (6.6 × 10^4^ parasites/ spleen and 2.1 × 10^6^ parasites/ liver) mono-infected mice exhibited lower parasite burdens in both organs compared to the 10:1 co-infected group at this time point (**Fig. 1C**, 1D and Supplementary Table S1). Collectively, these findings demonstrate that LS is capable of establishing a productive visceral infection independently in BALB/c mice, while co-infection with LD, particularly at the 10:1 co-inoculum ratio, exhibits maximal long-term parasite persistence.

ITS1 PCR followed by ImageJ-based densitometric analysis of the relative intensities of LD-and LS-specific PCR bands was used to estimate the LD: LS ratio in spleen and liver samples at different time points post-infection. During the early phase of infection (1.0-2.5 m.p.i.), the LD: LS ratio was variable, ranging from 1:1 to 1:10, with LS being equal to or more abundant than LD in several mice experiments (Supplementary Fig. S1 and Supplementary Table S2). At 3 m.p.i., ITS1 PCR performed on direct splenocytes and on “9-day” liver tissue cultures (which never transformed) revealed LD: LS ratios of 1:4 and 1:2 in the spleen, and 1:16 and 1:12 in the liver, following initial LD: LS co-inoculation ratios of 5:1 and 10:1, respectively (Supplementary Table S1).

In contrast, by 5 m.p.i., the LD: LS ratio had reversed in both spleen and liver. Direct spleen tissue showed ratios of 10:1 and 1.5:1, whereas liver tissue showed ratios of 1.5:1 and 10:1 in the 5:1 and 10:1 co-infection group, respectively (Supplementary Table S1). Similarly, analysis of spleen cultures maintained in M199 medium for 1–2 months (which never transformed) from mice sampled at 5-5.5 m.p.i. demonstrated that LD had become equal to or more abundant than LS, with LD: LS ratios ranging from 1:1 to 10:1 (Supplementary Fig. S2, S3 and Supplementary Table S3). Collectively, these findings suggest that although LS predominated during the early stages of co-infection, LD progressively recovered over time and became equal to or more abundant than LS at later stages of infection.

Quantitative PCR analysis at 3 and 5 m.p.i. confirmed the presence of parasite DNA in blood from all infection groups. At 3 m.p.i., LS DNA was detected in blood from mice infected with only LS confirming the presence of the parasite in circulation. Although LS parasite load (2.5 ×10^5^ gE/ mL) remained comparable to that of LD (2.5 ×10^5^ gE/ mL), a significantly higher circulating parasite burden was observed in the co-infection groups compared to the other infected groups. Parasite loads were estimated at (5.4 × 105 and 2.9 × 10^5^ gE/ mL) for 5:1 and 10:1 co-infections, respectively (**Fig. 1E** and Supplementary Table S1).

Interestingly, by 5 m.p.i., circulating parasite burden declined across all infection groups, with parasite loads ranging from (9.3 × 10^4^ to 2.01 × 10^5^ parasites/ mL). A significantly greater reduction (∼5.8 fold) in the 5:1 co-infection group (9.3 × 10^4^ gE/ mL) whereas only (∼1.7, 1.2 and 1.6 -fold) reduction was observed for LD (1.5 × 10^5^ gE /mL); LS (2 × 10^5^ gE/ mL) and 10:1 co-infection (1.8 × 10^5^ gE/ mL) respectively, compared to their corresponding 3 m.p.i loads (**Fig. 1F** and Supplementary Table S1). Collectively, these findings indicated that although circulating parasite burden decreased during chronic infection, parasites were still detected in the systemic circulation irrespective of the infection scenario.

#### 1.2. Lepsey NLV1 exhibits sustained persistence in murine tissues and circulation during LS mono- and co-infection

In preliminary experiments, RT-PCR analysis demonstrated that spleen cultures derived from LS mono-infected mice were positive for Lepsey NLV1 between 2 and 3 m.p.i. **(Supplementary Fig. S4, Supplementary Fig. S5).** In contrast to LS mono-infection, spleen cultures from co-infected mice remained Lepsey NLV1-positive over a broader period, from 1 to 5.5 m.p.i. **(Supplementary Fig. S6, Supplementary Fig. S7).** Although Lepsey NLV1 was readily detected following *ex vivo* culture, viral RNA was not detected directly in RNA later-preserved spleen tissues in these initial experiments **(Supplementary Fig. S7).** Analysis of peripheral blood revealed a distinct pattern: blood samples from co-infected mice were generally tested virus-positive, whereas those from LS mono-infected mice remained virus-negative. Specifically, Lepsey NLV1 was detected in blood as early as 1 m.p.i. in the 2:1 co-infection group **(Supplementary Fig. S8)** and persisted until at least 5 m.p.i. in the 10:1 co-infection group **(Supplementary Fig. S9).** Collectively, these preliminary observations suggested enhanced detectability and prolonged persistence of Lepsey NLV1 in co-infected animals relative to LS mono-infection. Representative samples found positive for virus, were re-confirmed by sequencing of Lepsey NLV1 **(Supplementary Table S4).**

Quantitative RT-PCR analysis was performed to estimate Lepsey NLV1 RNA levels in spleen and liver tissues at 3- and 5-m.p.i. At 3 m.p.i., Lepsey NLV1 was detected in the spleen tissues of LS mono-infected mice at (4.8 × 10^7^ gE/ spleen), while decreased virus titres were detected in the AG83 (2.5 × 10^7^ gE/ spleen) and 10:1 co-infection (3.1 × 10^7^ gE/ spleen) groups, respectively. In the liver tissues, virus titres remained comparable and were estimated at (8.6 × 10^7^, 9.8 × 10^7^ and 1 × 10^8^ gE/ liver) in the LS mono-infected, virus-positive AG83 and 10:1 co-infection group, respectively (**Fig 2A**, 2B, Supplementary Table S1).

**Figure 2.**
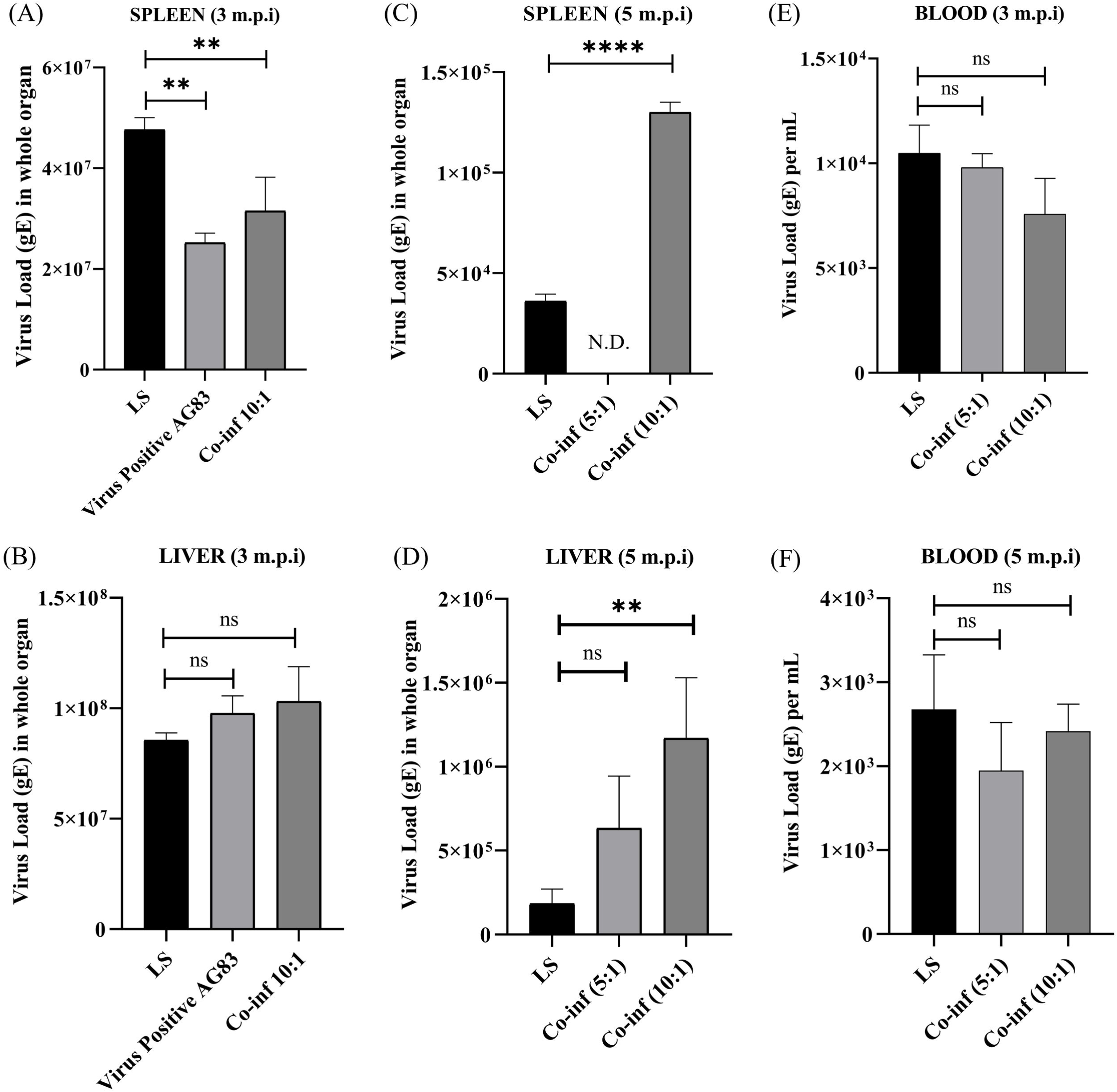
Temporal dynamics of Lepsey NLV1 burden in parasite infected BALB/c mice. Lepsey NLV1 burden was quantified by qRT-PCR in spleen, liver, and blood samples collected from BALB/c mice infected with LS, virus-positive AG83 laboratory isolate, or LD-LS co-infections at the indicated post-infection (p.i.) time points. **(A)** Virus load in spleen at 3 m.p.i **(B)** Virus load in liver at 3 m.p.i. **(C)** Virus load in spleen at 5 m.p.i. **(D)** Virus load in liver at 5 m.p.i **(E)** Virus load in blood at 3 m.p.i. **(F)** Virus load in blood at 5 m.p.i. Virus genome copies are expressed as genome equivalents (gE). The data were represented as mean ± SD of a minimum of three independent experiments. **N.D.** indicates that viral RNA was not detected. Statistical significance was determined using one-way ANOVA followed by Dunnett test where significance is denoted as p < 0.05 (*), p < 0.01 (**), p < 0.001 (***), p < 0.0001 (****), ns = not significant.

At 5 m.p.i., viral titres declined markedly in both organs. In the spleen tissues, Lepsey NLV1 was detected at (3.6 × 10^4^ gE/ spleen) in LS mono-infected group. Significantly greater virus load (1.3 × 10^5^ gE/ spleen) was detected in the 10:1 co-infection group compared to LS mono-infection at this time point. Notably, the 5:1 co-infection group was below the detection limit (possibly due to technical reasons). In contrast, liver tissues remained positive for Lepsey NLV1 in all LS-containing infection groups, with comparable viral titres of (1.9 × 10^5^ and 6.4 × 10^5^ gE/ liver) between LS-mono-infection and 5:1 co-infection group, respectively. Consistent with the virus titres in spleen tissues, significantly higher titre (1.2 × 10^6^ gE/ liver) was observed in the 10:1 co-infection compared to LS mono-infection group (**Fig 2C**, 2D and Supplementary Table S1). Collectively, these findings indicated a progressive decline in tissue viral burden during chronic infection over time, while demonstrating sustained viral persistence in LS-containing infections, particularly in mice receiving the 10:1 LD: LS co-infection.

A better estimate of viral load was obtained from subsequent experiments. Quantitative RT-PCR was done to analyse the blood samples (3 and 5 m.p.i.) from mice infected with LS, 5:1 or 10:1 co-culture of LD and LS. Viral RNA was detected in both LS mono-infected and co-infected mice at both time points. At 3 m.p.i., Lepsey NLV1 titres in blood were estimated at (1.0 × 10^4^, 9.7 × 10^3^ and 7.5 × 10^3^ gE/ mL) in LS mono-infection, 5:1 and 10:1 co-infection group, respectively, indicating comparable circulating viral titres across all LS-containing infection groups. By 5 m.p.i., viral titres declined across all groups. Virus titres decreased 3.8-fold (2.6 × 10^3^ gE/ mL), 5.1-fold (1.9 × 10^3^ gE/ mL) and 3.1-fold (2.4 × 10^3^ gE/ mL) in LS, 5:1 and 10:1 co-infection groups, respectively compared to their corresponding titres at 3 m.p.i. Notably, although absolute viral loads decreased over time, Lepsey NLV1 remained detectable in blood up to 5 m.p.i., indicating sustained systemic persistence during chronic infection (**Fig 2E**, 2F and Supplementary Table S1).

### 2. Microscopic visualization of cultured spleen tissue smears and peritoneal macrophages from mice infected with LD, LS and their co-infections

#### 2.1. Photomicrographs of *in vitro* cultured splenic stamp smears from parasite infected mice

Giemsa-stained smears were prepared post 14-days from *in vitro* culture of splenic tissues. The spleen was originally collected from infected animals at 5 m.p.i. and splenic tissues were subsequently maintained *in vitro* in Schneider’s transformation medium for 14 days. These smears revealed a successful morphological transformation of the parasites into their respective promastigote forms. Importantly, alongside the expected differentiation of LD, we provide the novel evidence that LS can persist within splenic tissue and subsequently transform, undergoing a clear transition from amastigote to promastigote stages under *in vitro* culture conditions **(Fig. 3A-C).** Histological stamp smears of visceral organs from 7 m.p.i. mice further corroborated our hypothesis **(Supplementary Fig. S10).**

**Figure 3.**
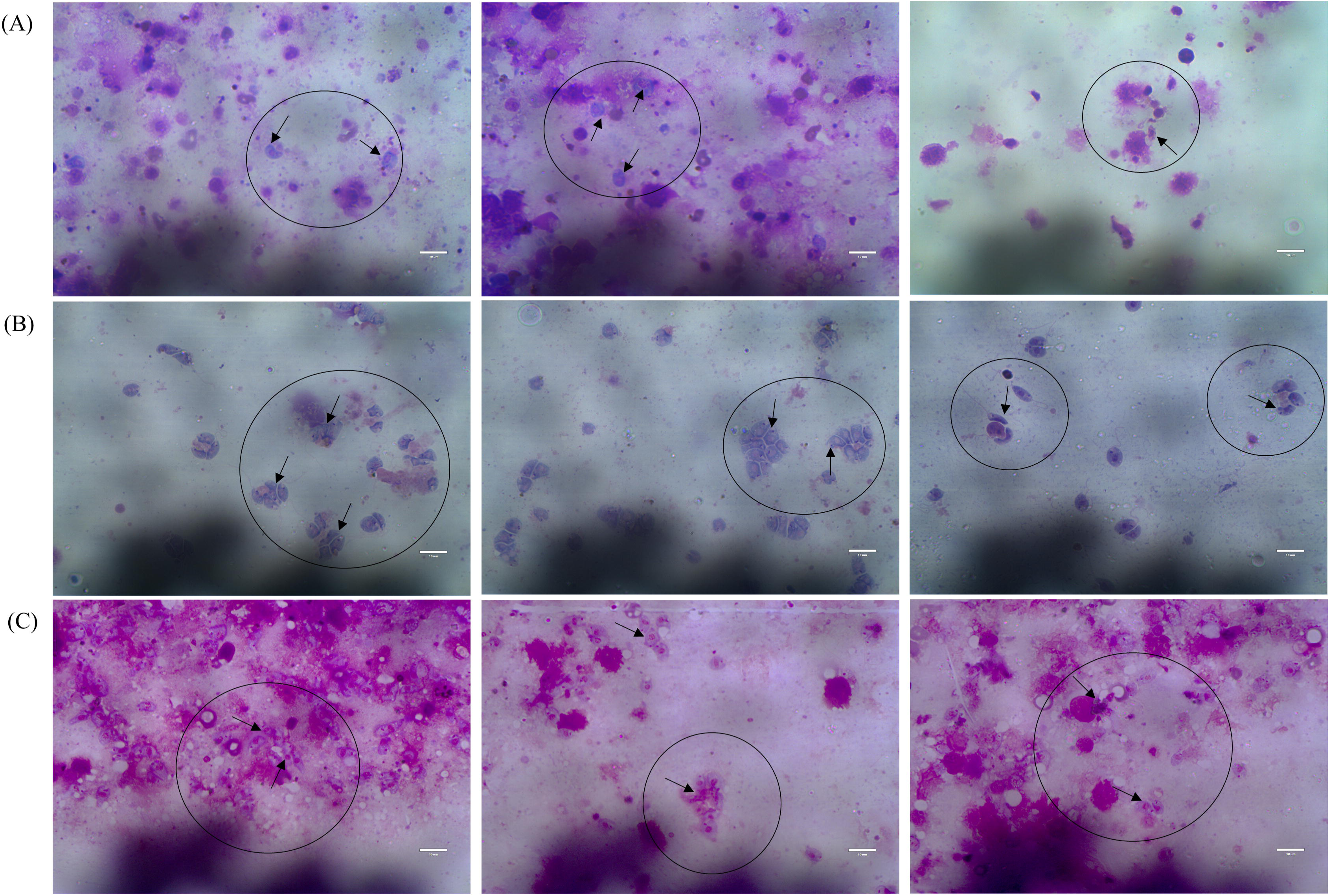
Representative Giemsa-stained histological stamp smears of spleen cultures recovered from infected BALB/c mice. Representative microscopic images (1000X magnification) of Giemsa-stained histological stamp smears prepared from 14-day *in vitro* cultured spleen samples recovered from BALB/c mice at 5 m.p.i. **(A)** LD infected mice; **(B)** LS infected mice; and **(C)** mice co-infected with LD and LS at a ratio of 10:1. Both LD and LS parasites were detected in the cultured spleen smears, exhibiting morphological differentiation from intracellular amastigote-like forms to extracellular promastigote forms during *in vitro* cultivation. Scale bar = 10 μm.

### 2.2. Murine peritoneal macrophage infection

Spherical-shaped amastigote images were observed under microscope at both 48 and 72 h.p.i. inside Giemsa-stained murine peritoneal macrophages. LS amastigotes, appearing as small peripheral dots within the infected cells, were detectable inside peritoneal macrophages, although they were less prominent compared to morphologically distinct LD amastigotes. We also observed apoptosis like features such as cell shrinkage and membrane blebbing in only LS-infected peritoneal macrophages **(Fig. 4).** Such features were morphologically absent in LD or co-culture infected cells. This is consistent with the fact that LD is known to prevent apoptosis (28,29).

**Figure 4.**
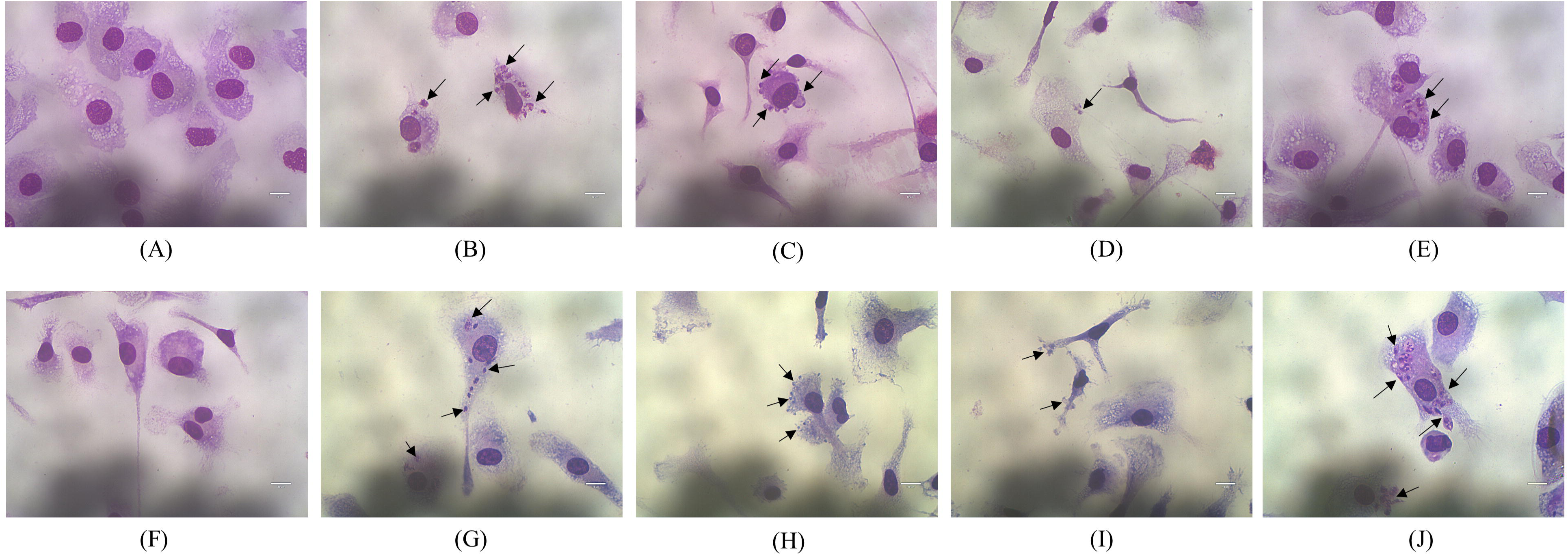
Representative Giemsa-stained images of parasite infected murine peritoneal macrophages. Representative Giemsa-stained microscopic images (1000X magnification) of murine peritoneal macrophages infected with LD, LS at multiplicities of infection (MOI) of 10:1 or 20:1, or with 10:1 co-infection of (LD: LS) respectively, followed by incubation for 48 h **(A–E)** and 72 h **(F–J)** post-infection. Representative fields illustrate intracellular parasite burden and morphological features under each infection condition. Scale bar = 10 μm.

**Fig. 5:**
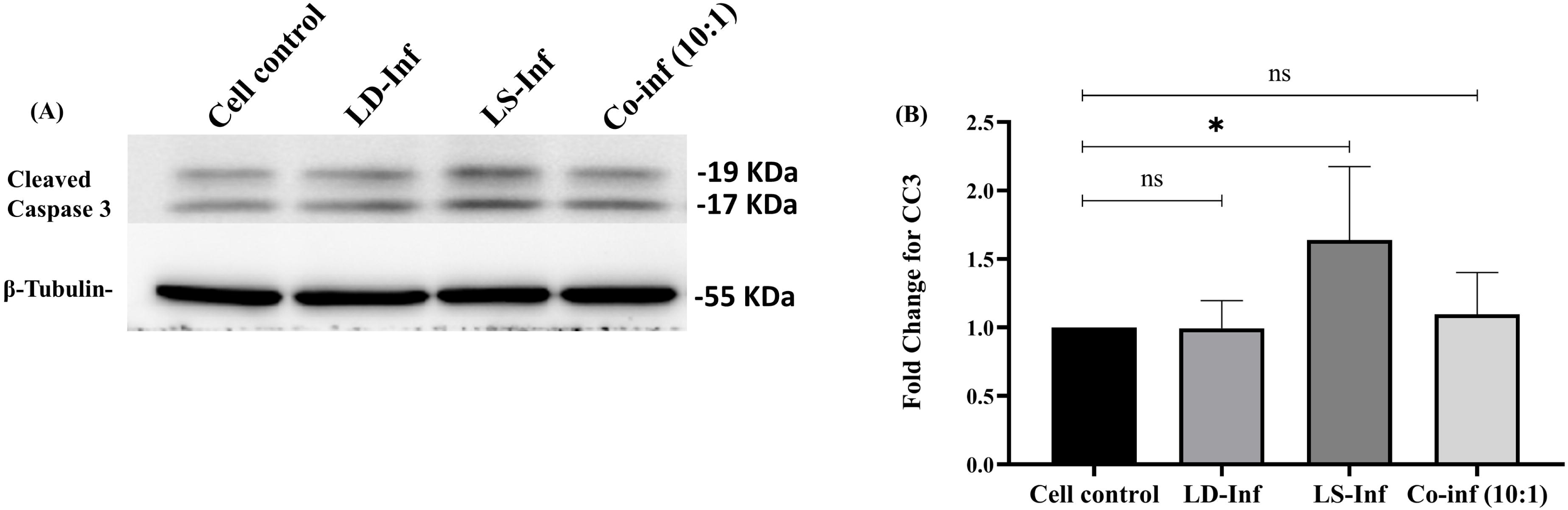
Western blot analysis of cleaved Caspase 3 expression for infected RAW264.7 cells at 72h.p.i. Immunoblot showing the expression of **(A)** cleaved Caspase 3 (upper panel) in infected RAW 264.7 cells at 72 h.p.i along with the corresponding internal control (lower panel). **(B)** Quantitative analysis was done by Image lab software (Bio-Rad) Statistical comparisons were made using one-way Anova followed by Dunnett test where ns-not significance, p < 0.05 (*).

### 3. LS infection induces an upregulation of cleaved caspase-3 expression, indicating activation of the apoptotic pathway

To further evaluate the apoptotic nature of LS, an *in vitro* western blot assay was performed using murine macrophage cells (RAW 264.7). At 72 h.p.i., LS-infected cells exhibited the highest level of cleaved caspase-3 (CC3) expression, a key marker of apoptosis, among the experimental groups. Densitometric analysis revealed that CC3 expression in LS-infected cells was significantly increased by approximately 1.6-fold compared to the un-infected cell control **(Fig: 5A-B).**

### 4. Reactivity of the mouse anti-LD and anti-LS immune serum against in vitro infected macrophages

Indirect immunofluorescence (IF) assays performed using mouse anti-LD and anti-LS serums yielded positive signals in *in vitro* infected macrophages. Notably, both anti-serums exhibited cross-reactivity, effectively staining macrophages infected with either LD or LS parasites. LD-infected RAW

264.7 macrophages displayed distinct fluorescence patterns, most prominently characterized by intense, diffuse cytoplasmic fluorescence, indicative of intracellular LD amastigotes. In comparison, LS-infected macrophages frequently exhibited a speckled fluorescence pattern, with multiple LS amastigotes detectable on the inner macrophage cell membrane as discrete fluorescent bodies. No reactivity was observed in control (uninfected) macrophages upon exposure to the respective immune serum **(Fig. 6).**

**Figure 6.**
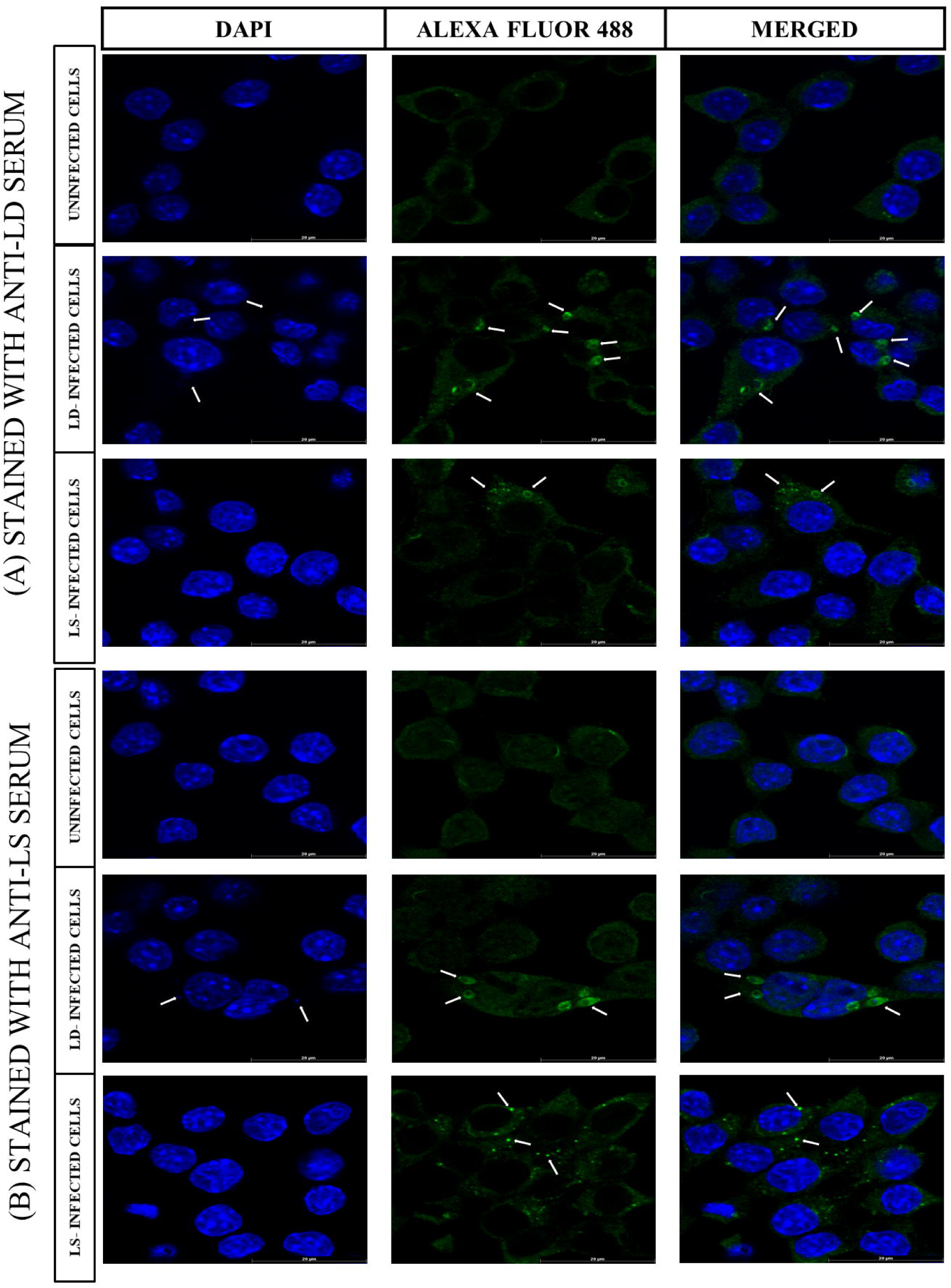
Confocal immunofluorescence microscopy demonstrating intracellular localization of parasites in infected macrophages. Representative confocal laser scanning microscopy images showing intracellular localization of parasites, LD and LS in infected macrophages under the indicated experimental conditions. Cells were fixed and immune-stained using **(A)** anti-LD or **(B)** anti-LS immune serum as the primary antibody, followed by a fluorescein (FITC)-conjugated secondary antibody to detect parasites (green). Host cell nuclei and parasite nuclear/ kinetoplast DNA were counterstained with DAPI (blue). Merged images demonstrate the intracellular localization of immunolabelled parasites. White arrows indicate representative intracellular LD or LS parasites. Scale bar = 20 μm.

### 5. Co-infection suppresses systemic IL-12 responses during disease progression

IL-12 (p40) ELISA performed with serum samples indicated that at 1 m.p.i., LD-infected mice showed significantly upregulated IL-12 expression by approximately 3.3-fold compared to the controls, whereas LS infected and co-culture (10:1)-infected mice showed significantly reduced IL-12 expression by approximately 1.2-fold and 1.4-fold, respectively compared to the LD-infected mice. A similar trend was observed at 3 m.p.i., LD-infected mice had the highest IL-12 levels in serum, indicating a strong Th1 response. Although LS-infected mice exhibited similar level compared to LD infection, co-infected mice (LD: LS, 10:1) displayed significantly reduced IL-12 by approximately 2.8-fold and 2.2-fold lower than LD- and LS-infected mice, respectively suggesting Th1 suppression. By 5 m.p.i., although IL-12 levels declined in LD-infected mice relative to its 3-months’ time point, LS-infected mice maintained a steady profile. Co-infected mice continued to show the lowest IL-12 levels by 1.4-fold and 1.5-fold lower than LD and LS groups, respectively **(Fig. 7).**

**Figure 7.**
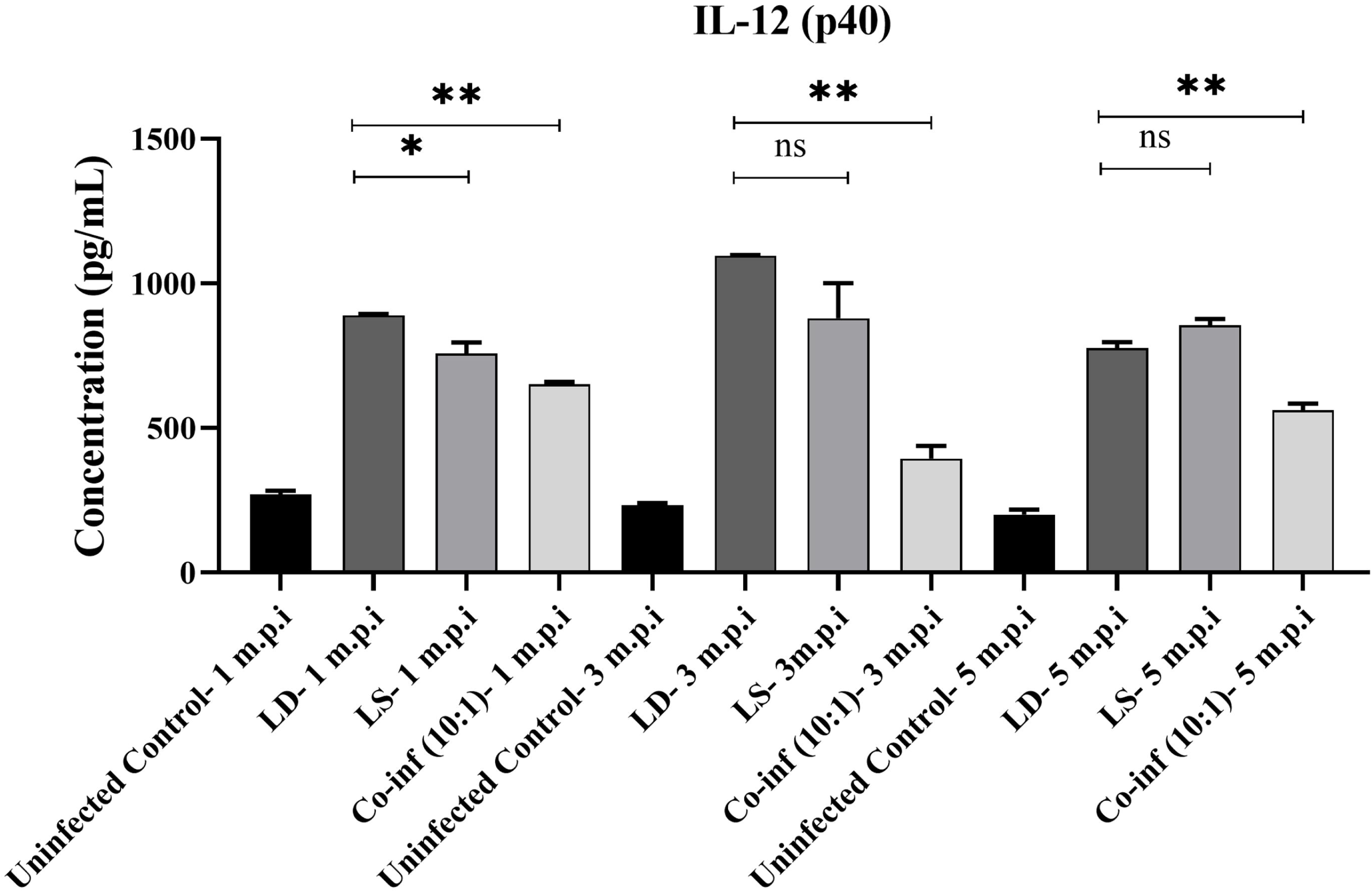
Serum IL-12 (p40) levels in mice following infection with LD, LS or 10:1 Co-infection. Serum IL-12 (p40) concentrations were quantified by ELISA at 1, 3, and 5 m.p.i. from uninfected control, LD, LS and 10:1 Co-infected mice. Data are presented as mean ± SD. Statistical comparisons were made using one-way Anova followed by Dunnett test where ns- not significant, p < 0.05 (*), p < 0.01 (**).

## Discussion

The present study provides an *in vivo* characterization of a tripartite infection system involving LD, LS and its viral endosymbiont Lepsey NLV1, revealing dynamic host-parasite-virus interactions that challenge the conventional view of visceral leishmaniasis (VL) as a mono-parasitic disease. By integrating longitudinal analyses of parasite burden, viral persistence and microscopic observations, we demonstrate that Lepsey NLV1-positive LS establishes sustained visceral infection in BALB/c mice both independently and during co-infection with LD. These findings uncover a temporally regulated shift in parasite dominance, prolonged systemic persistence of the viral endosymbiont and the unexpected capacity of a traditionally insect-restricted monoxenous trypanosomatid to adapt and persist within a mammalian host. Although LS has historically been considered non-pathogenic to mammals, its repeated detection in clinical VL and PKDL cases has raised questions regarding its biological significance (9–11). Our findings substantially strengthen this emerging concept by demonstrating prolonged persistence of LS in blood up to 5 m.p.i. and visceral organs up to 7 m.p.i., together with sustained detection of Lepsey NLV1. These observations extend our previous *in vitro* study demonstrating intracellular survival and replication of LS within macrophages and indicate that the parasite is capable of maintaining long-term infection under physiological conditions rather than representing transient contamination or passive carriage (12).

A key finding of this study is the temporal restructuring of parasite dominance during co-infection, where not only LD but also LS with its viral symbiont remained dominant parasite. Parasite quantifications revealed that by 3 m.p.i., spleen and liver parasite burdens had increased compared to the initial inoculum (2 × 10^7^ parasites/ mouse), indicating successful establishment and proliferation. Although a decline in parasite load was noted by 5 m.p.i. in mono-infections, 10:1 co-infected group particularly maintained more stable parasite levels, suggesting synergistic interactions that appeared to enhance persistence or immune evasion.

In case of 10:1 co-infection, early infection stages (1-3 m.p.i.) were characterised by a relative dominance of LS, particular in visceral organs, possibly due to enhanced replication or immune evasion aided by Lepsey NLV1. This observation aligns with emerging reports suggesting that LS is capable of infecting mammalian system and intracellular survival under permissive conditions (7,12). However, by 5 m.p.i., this trend reversed where LD regained dominance in visceral organs, indicating a competitive advantage in long-term persistence. This bi-phasic pattern was consistently observed over several *in vivo* experiments suggesting that LS may play a role in early niche establishment or immune modulation, potentially facilitating long term LD persistence. Supporting this interpretation, clinical studies have reported the sustained co-persistence of both parasites. Notably, LD and LS DNA remain detectable in PKDL skin biopsy samples even up to 7 years after apparent cure of visceral leishmaniasis, underscoring the long-term stability and clinical relevance of such co-infections (6).

The analysis of circulating parasite burden reveals a modest decline in overall parasitaemia between 3- and 5 m.p.i., suggesting partial host control during the transition from early to chronic infection. However, this apparent reduction in total parasite load contrasts sharply with the progressive dominance of LS in circulation over time. In co-infection scenarios, despite an initial inoculum strongly biased toward LD, the LD: LS ratio shifts dramatically in blood from 1:50 at 3 months to as high as 1:75 and 1:100 at 5 months, indicating that LS is not only persisting but also getting extra-viscerally disseminated.

One important observation from this study is the sustained Lepsey NLV1 persistence *in vivo*, particularly in co-infected animals. While LS-infected mice were virus-positive in spleen and blood from 2 to 3 m.p.i., virus was undetectable directly from primary tissues in earlier experiments unless amplified through culture, indicating low titres or compartmentalisation. Notably, viral loads were higher at earlier time points (3 m.p.i.) and remained detectable across visceral tissues and blood, with co-infection mostly enhancing viral detectability compared to LS mono-infection. This observation strongly supports the hypothesis that LD creates a permissive intracellular environment that enhances long-term virus persistence even in circulation. The load declined thereafter, yet remained within detectable ranges in LS and co-infection samples even at 5 m.p.i. This implied a temporally regulated control or adaptation of the host immune system to the viral presence. Importantly, the viral persistence in splenocytes and liver indicated that these organs might serve as virus reservoirs during chronic infection.

Importantly, this study provided direct morphological and experimental evidence for LS persistence within mammalian tissues, including its ability to differentiate from amastigote-like forms to promastigotes upon *in vitro* culture. The morphological transition of parasites in spleen culture not only supported the viability of LS within the mammalian host environment but also reinforced our hypothesis regarding its ability to establish and sustain infection. This finding challenges the long-standing dogma that LS is strictly monoxenous and incapable of sustaining infection in vertebrate hosts.

Microscopic analysis of infected murine peritoneal macrophages revealed that both LD and LS could survive within host macrophages, with LD showing more prominent intracellular presence. A preceding study using a similar methodology demonstrated microscopic visualization of *Leptomonas costoris* alongside LD in infected peritoneal macrophages from Syrian golden hamsters (30). The observation of LS amastigote-like forms within macrophages, albeit less prominent than LD, further supports its adaptive plasticity. Interestingly, LS infection was associated with apoptosis-like features in macrophages, in contrast to LD, which is known to inhibit host cell apoptosis (28,29).

The significant higher cleaved caspase-3 (CC3) expression in LS-infected murine macrophages further indicated that LS induces a stronger apoptotic response than LD infection. Since CC3 is a key marker of apoptosis, its elevated expression suggests enhanced activation of the apoptotic machinery following LS infection. In contrast, the relatively lower CC3 expression in LD-infected cells may indicate that LD suppresses or limits apoptotic signalling, potentially favouring intracellular parasite survival. Consistent with previous MTT observations, LS infection was associated with pronounced loss of macrophage viability, whereas increasing LD proportions during co-infection progressively preserved host-cell viability (12). Overall, these findings suggested fundamentally different host-pathogen interaction strategies, where LS may induce more pro-apoptotic intracellular environment potentially aiding parasite dissemination or immune evasion.

The immunofluorescence analysis using mouse-derived anti-LD and anti-LS serums provides important insight into the host immune response generated during infection. Notably, the successful detection of intracellular parasites using anti-LS serums confirms that LS is capable of eliciting a robust humoral immune response, despite its traditional classification as a monoxenous, insect-restricted organism. This observation strongly supports the notion that LS not only survives within the mammalian host but is also immunologically recognized, leading to the production of specific antibodies. The observed cross-reactivity between anti-LD and anti-LS serum further suggests the presence of shared or conserved antigenic determinants between the two species, reflecting evolutionary or structural similarities in surface or intracellular proteins. Collectively, these findings reinforce the concept that LS actively engages with the host immune system and is not merely a passive co-infecting organism, thereby underscoring its potential role in shaping host-parasite immune dynamics during co-infection.

The differential IL-12 responses suggest that LD and LS distinctly modulate host immunity during infection. The pronounced elevation of IL-12 in LD-infected mice, particularly during the early and intermediate stages, indicates a strong Th1-associated response, which is important for IFN-γ-mediated macrophage activation and parasite control. In contrast, the consistently reduced IL-12 levels in co-infected mice suggest that the presence of LS suppresses the Th1 response induced by LD. These findings are consistent with our previously published observations, which demonstrated reduced IL-18 levels in human serum samples from co-infected cases compared with those from LD-only infections (15). Collectively, these findings suggest that LS may modulate the LD-induced inflammatory response, creating a comparatively immunosuppressive environment during mixed infection that could potentially favour parasite persistence.

Future investigations should focus on deciphering the immunological and molecular mechanisms underlying this tripartite interaction, particularly the role of innate immune signalling pathways (e.g., TLR3, inflammasome activation) in response to viral RNA. The potential of LS and its virus to modulate treatment response, disease severity, or relapse also warrants exploration, especially in light of analogous findings with LRV in cutaneous leishmaniasis (16,18, 31–33). In conclusion, this study establishes a novel paradigm of multi-pathogen and virus-associated parasitism in VL, demonstrating that interactions between dixenous parasites, and viral endosymbionts can significantly influence infection dynamics. These findings not only expand our understanding of VL pathogenesis but also highlight the need to reconsider co-infection and virome contributions in parasitic diseases, with potential implications for diagnosis, prognosis, and therapeutic intervention.

## Supporting information

with all infection groups showing greater parasite accumulation in the liver than in the spleen (Fig. 1A, 1B and Supplementary Table S1).

## ETHICS STATEMENT

All experimental protocols were performed in accordance with the Committee for the Control and Supervision of Experiments on Animals (CCSEA) guidelines. The study protocol was approved by Institutional Animal Ethics Committee (IAEC, CSIR-IICB, Kolkata). The experiments on animals comply to the ARRIVE guidelines. Approval identifier: IICB/AEC/Meeting/Sep/2023/6.

## AUTHORS’ CONTRIBUTION

S.B., S.D. and P.D.S. conceived and designed the experiments. S.D., P.D.S. performed the experiments equally. R.C. also performed many of the experiments with S.D. and P.D.S. S.D. and P.D.S. performed data analysis and S.B. performed critical analysis of the data. S.B. provided funding, reagents, materials and analysis tools. S.B., S.D. and P.D.S. jointly wrote the original draft of the manuscript. All authors critically reviewed and modified and agreed on the current version of the manuscript.

## ACKNOWLEDGEMENT

Authors acknowledge the support received from the Director of CSIR IICB. S.B. also acknowledges AcSIR and ICMR for support. S.D. and P.D.S acknowledges the support of UGC for their UGC-Research Fellowships. R.C. acknowledges ICMR for support as Project Research Scientist-II. The authors would also like to acknowledge the use of relevant facilities at CSIR-IICB including the Central Instrumentation Facility (CIF). The authors acknowledge CSIR-IICB for providing all laboratory facilities for conducting the current work.

## DATA AVAILABILITY

All data have been provided in main manuscript and supplementary document. Further information and resource requests should be directed to and will be fulfilled by the corresponding author, Dr. Subhajit Biswas.

## Supporting Information Legends

**Table S1. Quantification of parasite burden and virus load in BALB/c mice following mono and co-infection.**

Parasite burden and *Leptomonas seymouri* narna-like virus 1 (Lepsey NLV1) load were determined in the spleen, liver, and blood of BALB/c mice at 3- and 5-months post-infection (m.p.i.) following infection with *Leishmania donovani* (LD), *Leptomonas seymouri* (LS), virus-positive AG83 or LD-LS co-infections (5:1 and 10:1) at 2 × 10^7^ parasites/ mouse. Parasite burden was quantified by species-specific ITS1 qPCR, while viral load was measured by Lepsey NLV1-specific qRT-PCR. For co-infected samples, individual LD and LS parasite burdens are presented along with the corresponding LD: LS ratios. “Not detected” indicates that Lepsey NLV1 RNA was below the assay detection limit. Data are presented as mean ± SD from a minimum of three independent experiments.

**Figure S1. Representative image of ITS1 PCR analysis from spleen culture samples of BALB/c mice infected with parasites.**

M: marker; L1: represents DNA from spleen cultures of mice infected with only LD after 1-m.p.i. and 13 days culture; L2: represents 10:1 co-culture infected sample after 1-m.p.i. and 13 days culture; L3: represents 2:1 co-culture infected sample after 1-m.p.i. and 5 days culture; L4-L5: represents DNA from spleen cultures of mice infected with only LS after 2-m.p.i. followed by 1 and 2 months culture, respectively (not relevant to this image); L6-L7: represents DNA from spleen cultures of mice infected with AG83 lab isolate after 2.5-m.p.i. followed by 1 and 2-months culture respectively; L8 and L9: positive control for LD and LS respectively, L10: negative control, (where L represents Lane number); L2 white arrow indicates the presence and detection of both LS and LD positive bands in 13 days spleen culture at 1 m.p.i. for 10:1 co-culture infected mouse.

**Table S2. Estimated parasite copy number and (LD: LS) ratio in parasites recovered from infected BALB/c mice following visceral organ culture.**

Spleen cultures established from BALB/c mice infected with LD, LD: LS co-infections (2:1 or 10:1), or AG83 laboratory isolate were analysed for parasite burden after the indicated duration of *in vivo* infection and subsequent *ex vivo* culture. Estimated parasite copy numbers were determined by ITS1 PCR followed by ImageJ analysis and are presented as approximate ranges. Samples SS-1, SS-2, and SS-3 represent independent spleen culture isolates obtained after 1 m.p.i, whereas AG83 samples were recovered after 2.5 m.p.i and analysed following 1 or 2 months of culture.

**Figure S2: Representative image of ITS1 PCR analysis from spleen culture samples of BALB/c mice infected with only LS; or 5:1; 10:1 co-cultures of LD: LS or AG83 lab isolate.**

M: Marker; L1-L2: represents DNA from spleen cultures of mice infected with only LS after 2 m.p.i. followed by 1-month and 2-months cultures, respectively. L3-L4: represents DNA from spleen cultures of mice infected with a co-culture (LD: LS) of 5:1 after 5 m.p.i. followed by 1-month culture (representative of two different biological replicates). L5: represents a similar DNA sample from mice infected with co-culture (LD: LS) of 10:1 at 5 m.p.i. followed by 1-month culture. L6-L7: represents spleen DNA samples from mice infected with AG83 lab isolate after 5.5 m.p.i. followed by 1-month culture (representative of two different biological replicates). L8-L9: represents positive control DNA from LD and LS, respectively; L10: represents negative control (where L represents lane number); white arrow (in L3-L4 and L6-L7) indicates the presence and detection of both LS and LD positive bands in 1-month spleen cultures at 5 months and 5.5 months for a 5:1 co-culture and virus-positive AG83 lab isolate infected mice respectively.

**Figure S3: Representative image of ITS1 PCR analysis from the DNA samples after ethanol precipitation of DNA.**

M: Marker; L1-L2: represents DNA from spleen cultures of mice infected with only LS after 2 m.p.i. followed by 1-month and 2-months cultures, respectively. L3: represents the same co-culture 10:1 infected spleen DNA sample, which was repeated with maximum template (not relevant). L4-L5: represents positive control DNA from LD and LS, respectively; L6: represents negative control (where L represents lane number), L1-L2 white arrow indicates the presence and detection of LS positive band in 1- and 2-months spleen culture at 2 m.p.i. for a LS infected mouse respectively.

**Table S3: Estimated parasite copy number and (LD: LS) ratio in parasites recovered from infected BALB/c mice following visceral organ culture.**

Spleen cultures established from BALB/c mice infected with LS, LD: LS co-infections (5:1 or 10:1), or AG83 laboratory isolate were analysed for parasite burden after the indicated duration of *in vivo* infection and subsequent *ex vivo* culture. Estimated parasite copy numbers were determined by ITS1 PCR followed by ImageJ analysis and are presented as approximate ranges.

**Figure S4: Representative image of LS DNA PCR and RT-PCR analysis from spleen culture samples of BALB/c mice infected with only LS at 2 m.p.i. followed by a 1-month culture.**

M: Marker; L1: represents spleen DNA; L2: represents positive control DNA from LS; L3: represents negative control. L4-L6: first round RT-PCR products from spleen RNA, positive control for Lepsey NLV1 and negative control respectively; L7-L9: represents second round PCR products from spleen RNA, positive control for Lepsey NLV1 and negative control respectively (where L represents lane number); L7 white arrow indicates the presence and detection of virus positive band in 1-month spleen culture at 2 m.p.i. for a LS infected mouse.

**Figure S5: Representative image of RT-PCR analysis for the detection of Lepsey NLV1 in splenocyte samples from infected BALB/c mice after 3 m.p.i.**

M: marker; L1-L4: represents first round PCR products from LD, LS, (LD: LS) 5:1 and 10:1 co-infected sample, respectively; L5 and L6: represents first round PCR products of positive control and negative control respectively. L7-L10: represents the second round PCR products from LD, LS, 5:1 and 10:1 co-infected sample, respectively; L11-L12: Second round PCR samples for positive control and negative control respectively (where L represents Lane number); L8-L10 white arrow indicates the presence and detection of virus positive band at 3 m.p.i. from mice infected with LS, 5:1 and 10:1 co-infection respectively.

**Figure S6: Representative image of RT-PCR analysis of RNA from spleen samples of BALB/c mice infected with parasites.**

M: marker; L1: represents negative control for first round PCR; L2-L4: represents first round PCR products from spleen of mice infected with only LD after 1-m.p.i. followed by 13 days of tissue culture, (LD: LS) 10:1 co-infected spleen sample after 1-m.p.i. followed by 13 days of tissue culture; 2:1 co-infected spleen sample after 1-m.p.i. followed by 5 days of tissue culture respectively; L5: represents first round positive control from RNA from Lepsey NLV1; L6: second round PCR for negative control; L7-L9: represents second round PCR from same spleen of mice infected with only LD after 1-m.p.i. followed by 13 days of tissue culture, 10:1 co-infected spleen sample after 1-m.p.i. followed by 13 days of tissue culture; 2:1 co-infected spleen sample after 1-m.p.i. followed by 5 days of tissue culture respectively; L10: Second round PCR samples for positive control (where L represents Lane number). L8-L9 white arrow indicates the presence and detection of virus in 13 days and 5 days spleen tissue cultures at 1-m.p.i. for a 10:1 and 2:1 co-infected mouse.

**Figure S7: Representative image of RT-PCR analysis for the detection of Lepsey NLV1 from BALB/c mice spleen samples infected with AG83 lab isolate.**

M: marker; L1-L2: represents first round PCR products from spleen tissue stored in RNA Later after 5.5 m.p.i. (representative of two different biological replicates); L3-L4: represents first round PCR products from the same samples after 5.5 m.p.i. followed by a 1-month tissue culture (representative of two different biological replicates); L5-L6: represents the first round PCR products of the positive control and negative control, respectively. L7-L8: represents the second round PCR of the same spleen samples directly from tissue (representative of two different biological replicates); L9-L10: represents the second round PCR of the same spleen samples after 1 month of tissue culture (representative of two different biological replicates); L11-L12: Second round PCR samples for positive control and negative control respectively (where L represents Lane number). White arrow at L9-L10 indicates the detection and presence of virus in 1-month spleen tissue cultures at 5.5 m.p.i. for a virus-positive AG83 lab isolate infected mice.

**Figure S8: Representative image of RT-PCR analysis from blood samples of BALB/c mice infected with parasites.**

M: marker; L1-L4: represents first PCR products for negative control, RNA samples of spleen infected with LD, 10:1 and 2:1 co-infection of LD: LS respectively after 1-m.p.i. L5-L8: represents the second round PCR products of the RNA samples of spleen infected with LD, 10:1 and 2:1 co-infection of LD: LS after 1-m.p.i. and negative control respectively, (where L represents Lane number). L7 white arrow suggests the presence and detection of virus in blood at 1-m.p.i. for a 2:1 co-infected mouse.

**Figure S9: Representative image of RT-PCR analysis from blood samples of BALB/c mice infected with parasites.**

M: marker; L1 and L3: represents first round PCR products from blood infected with LS and (LD: LS) 10:1 co-infection after 2 m.p.i. respectively. L2 and L4: represents the first round PCR products of same samples after 5 m.p.i. respectively. L5 and L6: represents positive and negative control respectively. L7 and L9: represents second round PCR products from blood infected with LS and 10:1 co-infection after 2 m.p.i. respectively. L8 and L10: represents the second round PCR products of same samples after 5 m.p.i. respectively; L11: Positive control for Lepsey NLV1; L12: Negative control (where L represents Lane number). L10 white arrow indicates the presence and detection of virus in blood at 5-m.p.i. for a 10:1 co-infected mouse.

**Table S4. Multiple sequence alignment of the amplified Lepsey NLV1 fragment recovered from experimentally infected BALB/c mice with closely related Lepsey NLV1 sequences identified by BLAST analysis.**

The amplified Lepsey NLV1 sequences obtained from infected mouse samples were aligned with representative closely related Lepsey NLV1 sequences retrieved through BLAST analysis to assess sequence similarity. Sample 2L represents the spleen culture isolate obtained from a mouse infected with a 2:1 co-infection (LD: LS) and collected at 1 m.p.i. Sample L10 represents the spleen culture isolate from a mouse infected with a 10:1 co-infection (LD: LS) at 1 m.p.i. Sample 4TA1 represents a blood sample collected from a mouse infected with a 10:1 co-infection (LD: LS) at 5 m.p.i. Conserved nucleotides are indicated by alignment with homologous Lepsey NLV1 sequences, demonstrating high sequence identity between the experimentally recovered viral isolates and previously reported Lepsey NLV1 sequences.

**Figure S10: Representative microscopic views (1000X) of Giemsa-stained histological stamp smear of Spleen and Liver from mice at 7-m.p.i.**

Histological stamp smears of visceral organs (A) Spleen and (B) Liver from BALB/c mice infected with LD, LS, virus-positive AG83 and 10:1 co-infection of (LD: LS). Control mice were injected with sterile 1X PBS.

