## Supplementary material for "Tripartite host-parasite-virus interactions reshape chronic visceral leishmaniasis through persistent *Leptomonas seymouri* co-infection": with all infection groups showing greater parasite accumulation in the liver than in the spleen (Fig. 1A, 1B and Supplementary Table S1).

**Supplementary Data**

**Table S1. Quantification of parasite burden and virus load in BALB/c mice following mono and co-infection.** Parasite burden and *Leptomonas seymouri* narna-like virus 1 (Lepsey NLV1) load were determined in the spleen, liver, and blood of BALB/c mice at 3- and 5-months post-infection (m.p.i.) following infection with *Leishmania donovani* (LD), *Leptomonas seymouri* (LS), virus-positive AG83 or LD-LS co-infections (5:1 and 10:1) at 2 x $10^{7}$ parasites/mouse. Parasite burden was quantified by species-specific ITS1 qPCR, while viral load was measured by Lepsey NLV1-specific qRT-PCR. For co-infected samples, individual LD and LS parasite burdens are presented along with the corresponding LD: LS ratios. "Not detected" indicates that Lepsey NLV1 RNA was below the assay detection limit. Data are presented as mean ± SD from a minimum of three independent experiments.

| Sample Name | Parasite Load | | | Viral Load by qRT-PCR | Parasite: Virus Ratio | Fold increase w.r.t LS |
|---|---|---|---|---|---|---|
| | By qPCR | | Ratio (LD: LS) | | | |
| | Spleen – 3 m.p.i. | | | | | |
| LD-SPLEEN | 1.1 (± 0.1) x $10^{8}$ | | | Not Detected | | |
| LS- SPLEEN | 6.8 (± 0.4) x $10^{7}$ | | | 4.8 (± 0.2) x $10^{7}$ | 1: 0.7 | |
| CO-INF (5:1)-SPLEEN | 6.6 (± 1) x $10^{7}$ | 1.3 (± 0.2) x $10^{7}$ (LD) | 1:4 | Sample N.A. | - | - |
| | | 5.3 (± 0.8) x $10^{7}$ (LS) | | | | |
| AG83-SPLEEN | 4.5 (± 0.4) x 105 | 2.2 (± 0.2) x $10^{3}$ (LD) | 1:200 | 2.5 (± 0.2) x $10^{7}$ | 1: 55.6 | 79.4 |
| | | 4.5 (± 0.4) x $10^{5}$ (LS) | | | | |
| CO-INF (10:1)-SPLEEN | 7.3 (± 0.2) x $10^{7}$ | 2.4 (± 0.07) x $10^{7}$ (LD) | 1:2 | 3.1 (± 0.7) x $10^{7}$ | 1:0.6 | 0.9 |
| | | 4.9 (± 0.1) x $10^{7}$ (LS) | | | | |
| | Spleen- 5 m.p.i. | | | | | |
| LD-SPLEEN | 1.3 (± 0.04) x $10^{5}$ | | | Not Detected | | |

| | | | | | | |
|---|---|---|---|---|---|---|
| **LS- SPLEEN** | 6.6 (± 0.6) x $10^4$ | | | 3.6 (± 0.4) x $10^4$ | 1:0.5 | |
| **CO-INF (5:1)- SPLEEN** | 3.4 (± 0.2) x $10^5$ | 3.1 (± 0.2) x $10^5$ (LD)<br>3.1 (± 0.2) x $10^4$ (LS) | 10:1 | Not Detected | - | - |
| **CO-INF (10:1)- SPLEEN** | 5 (± 0.01) x $10^5$ | 3 (± 0.01) x $10^5$ (LD)<br>2 (± 0.01) x $10^5$ (LS) | 1.5:1 | 1.3 (± 0.05) x $10^5$ | 1:0.7 | 1.4 |
| | **Liver- 3 m.p.i.** | | | | | |
| **LD- LIVER** | 4.1 (± 0.1) x $10^9$ | | | Not Detected | | |
| **LS- LIVER** | 5.3 (± 0.4) x $10^9$ | | | 8.6 (± 0.3) x $10^7$ | 1:0.02 | |
| **CO-INF (5:1)- LIVER** | 6.1(± 0.3) x $10^9$ | 3.6 (± 0.2) x $10^8$ (LD)<br>5.7 (± 0.3) x $10^9$ (LS) | 1:16 | Sample N.A. | - | - |
| **AG83 - LIVER** | 1.6 (± 0.1) x $10^8$ | 1.6 x $10^4$ (LD)<br>1.6 (± 0.1) x $10^8$ (LS) | 1:10000 | 9.8 (± 0.8) x $10^7$ | 1:0.6 | 30 |
| **CO-INF (10:1)- LIVER** | 6.5(± 0.7) x $10^9$ | 5 (± 0.5) x $10^8$ (LD)<br>6 (± 0.6) x $10^9$ (LS) | 1:12 | 1.0 (± 0.2) x $10^8$ | 1:0.02 | 1 |
| | **Liver- 5 m.p.i.** | | | | | |
| **LD- LIVER** | 2.5 (± 0.1) x $10^6$ | | | Not Detected | | |
| **LS- LIVER** | 2.1 (± 0.2) x $10^6$ | | | 1.9 (± 0.9) x $10^5$ | 1:0.1 | |
| **CO-INF (5:1)- LIVER** | 2.4 (± 0.1) x $10^6$ | 1.4 (± 0.10) x $10^6$ (LD)<br>9.6 (± 0.4) x $10^5$ (LS) | 1.5:1 | 6.3 (± 3.1) x $10^6$ | 1:0.7 | 7 |
| **CO-INF (10:1)- LIVER** | 5.1 (± 0.2) x $10^6$ | 4.6 (± 0.1) x $10^6$ (LD)<br>4.6 (± 0.1) x $10^5$ (LS) | 10:1 | 1.2 (± 0.4) x $10^6$ | 1:2.6 | 26 |
| **Sample Name** | **Parasite Load (/ml)** | | | **Viral Load (/ml) by qRT-PCR** | | |
| | **By qPCR** | | **Ratio (LD: LS)** | | | |
| | **Blood- 3 m.p.i.** | | | | | |
| **LD- BLOOD** | 2.5 (± 0.1) x $10^5$ | | | Not Detected | | |

| | | | | | | |
|---|---|---|---|---|---|---|
| **LS- BLOOD** | 2.5 (± 0.05) x $10^5$ | | | 1 (± 0.1) x $10^4$ | 1: 0.04 | |
| **CO-INF (5:1)- BLOOD** | 5.4 (± 0.2) x $10^5$ | 1.0 (± 0.04) x $10^4$ (LD)<br>5.3 (± 0.04) x $10^5$ (LS) | 1:50 | 9.7 (± 0.7) x $10^3$ | 1: 0.02 | 0.5 |
| **CO-INF (10:1)- BLOOD** | 2.9 (± 0.1) x $10^5$ | 5.6 (± 0.2) x $10^3$ (LD)<br>2.8 (± 0.1) x $10^5$ (LS) | 1:50 | 7.5 (± 1.7) x $10^3$ | 1: 0.03 | 0.75 |
| | **Blood- 5 m.p.i.** | | | | | |
| **LD- BLOOD** | 1.5 (± 0.1) x $10^5$ | | | Not Detected | | |
| **LS- BLOOD** | 2 (± 0.03) x $10^5$ | | | 2.6 (± 0.7) x $10^3$ | 1: 0.01 | |
| **CO-INF (5:1)- BLOOD** | 9.3 (± 0.1) x $10^4$ | 1.2 (± 0.02) x $10^3$ (LD)<br>9.1 (± 0.1) x $10^4$ (LS) | 1:75 | 1.9 (± 0.6) x $10^3$ | 1: 0.02 | 2 |
| **CO-INF (10:1)- BLOOD** | 1.8 (± 0.1) x $10^5$ | 1.7 (± 0.1) x $10^3$ (LD)<br>1.7 (± 0.1) x $10^5$ (LS) | 1:100 | 2.4 (± 0.3) x $10^3$ | 1: 0.01 | 1 |

**Figure S1. Representative image of ITS1 PCR analysis from spleen culture samples of BALB/c mice infected with parasites.**

M: marker; L1: represents DNA from spleen cultures of mice infected with only LD after 1-m.p.i. and 13 days culture; L2: represents 10:1 co-culture infected sample after 1-m.p.i. and 13 days culture; L3: represents 2:1 co-culture infected sample after 1-m.p.i. and 5 days culture; L4-L5: represents DNA from spleen cultures of mice infected with only LS after 2-m.p.i. followed by 1 and 2 months culture, respectively (not relevant to this image); L6-L7: represents DNA from spleen cultures of mice infected with AG83 lab isolate after 2.5-m.p.i. followed by 1 and 2- months culture respectively; L8 and L9: positive control for LD and LS respectively, L10: negative control, (where L represents Lane number); L2 white arrow indicates the presence and detection of both LS and LD positive bands in 13 days spleen culture at 1 m.p.i. for 10:1 co-culture infected mouse.

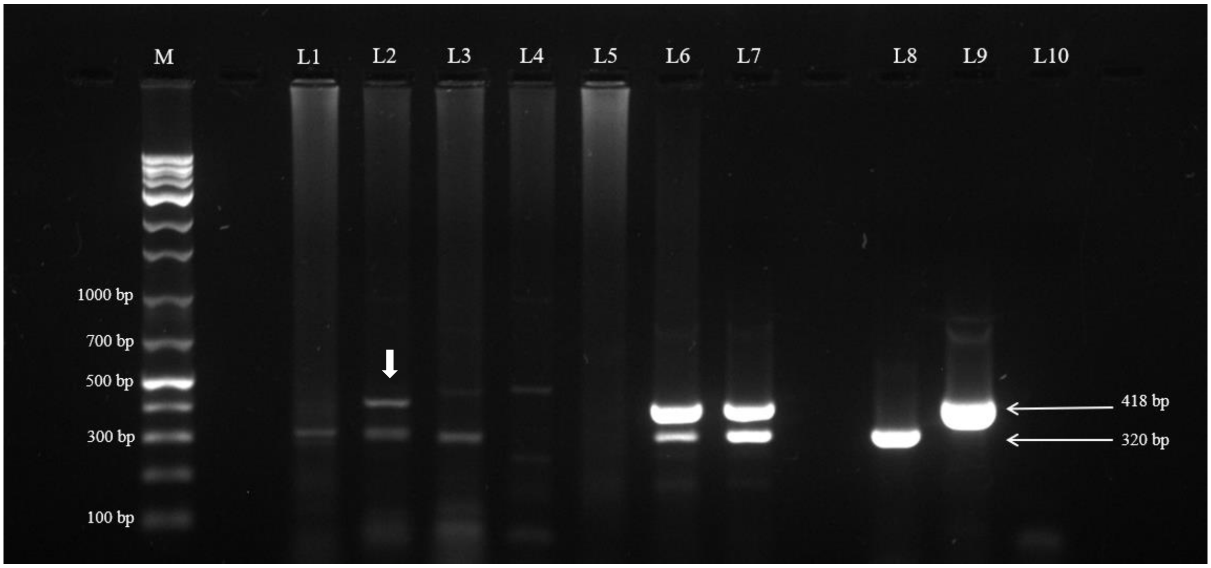

**Table S2. Estimated parasite copy number and (LD: LS) ratio in parasites recovered from infected BALB/c mice following visceral organ culture.**

Spleen cultures established from BALB/c mice infected with LD, LD: LS co-infections (2:1 or 10:1), or AG83 laboratory isolate were analysed for parasite burden after the indicated duration of *in vivo* infection and subsequent *ex vivo* culture. Estimated parasite copy numbers were determined by ITS1 PCR followed by ImageJ analysis and are presented as approximate ranges. Samples SS-1, SS-2, and SS-3 represent independent spleen culture isolates obtained after 1m.p.i, whereas AG83 samples were recovered after 2.5 m.p.i. and analysed following 1 or 2 months of culture.

| SAMPLE | INFECTION/CULTURE CONDITIONS | ESTIMATED COPY NUMBER | RATIO (LD: LS) |
|---|---|---|---|
| **(GROUP LD) SS-1 LD** | 1 m.p.i. / 13 days culture | $10^{3}$-$10^{4}$ | |
| **(GROUP 10:1) SS-2 LD** | 1 m.p.i. /13 days culture | $10^{3}$-$10^{4}$ | 1:1 |
| **(GROUP 10:1) SS-2 LS** | | $10^{3}$-$10^{4}$ | |
| **(GROUP 2:1) SS-3** | 1 m.p.i. / 5 days culture | $10^{3}$-$10^{4}$ | **Only LD detected** |
| **AG83 -LD** | 2.5 m.p.i. / 1-month culture | $10^{4}$ - $10^{5}$ | 1:10 |
| **AG83-LS** | | $10^{5}$- $10^{6}$ | |
| **AG83 - LD** | 2.5 m.p.i. / 2 months culture | $10^{6}$ -.> $10^{7}$ | 1:10 |
| **AG83- LS** | | .> $10^{7}$ | |

**Figure S2: Representative image of ITS1 PCR analysis from spleen culture samples of BALB/c mice infected with only LS; or 5:1; 10:1 co-cultures of LD: LS or AG83 lab isolate.**

M: Marker; L1-L2: represents DNA from spleen cultures of mice infected with only LS after 2 m.p.i. followed by 1-month and 2-months cultures, respectively. L3-L4: represents DNA from spleen cultures of mice infected with a co-culture (LD: LS) of 5:1 after 5 m.p.i. followed by 1-month culture (representative of two different biological replicates). L5: represents a similar DNA sample from mice infected with co-culture (LD: LS) of 10:1 at 5 m.p.i. followed by 1-month culture. L6-L7: represents spleen DNA samples from mice infected with AG83 lab isolate after 5.5 m.p.i. followed by 1-month culture (representative of two different biological replicates). L8-L9: represents positive control DNA from LD and LS, respectively; L10: represents negative control (where L represents lane number); white arrow (in L3-L4 and L6-L7) indicates the presence and detection of both LS and LD positive bands in 1-month spleen cultures at 5 months and 5.5 months for a 5:1 co-culture and virus-positive AG83 lab isolate infected mice respectively.

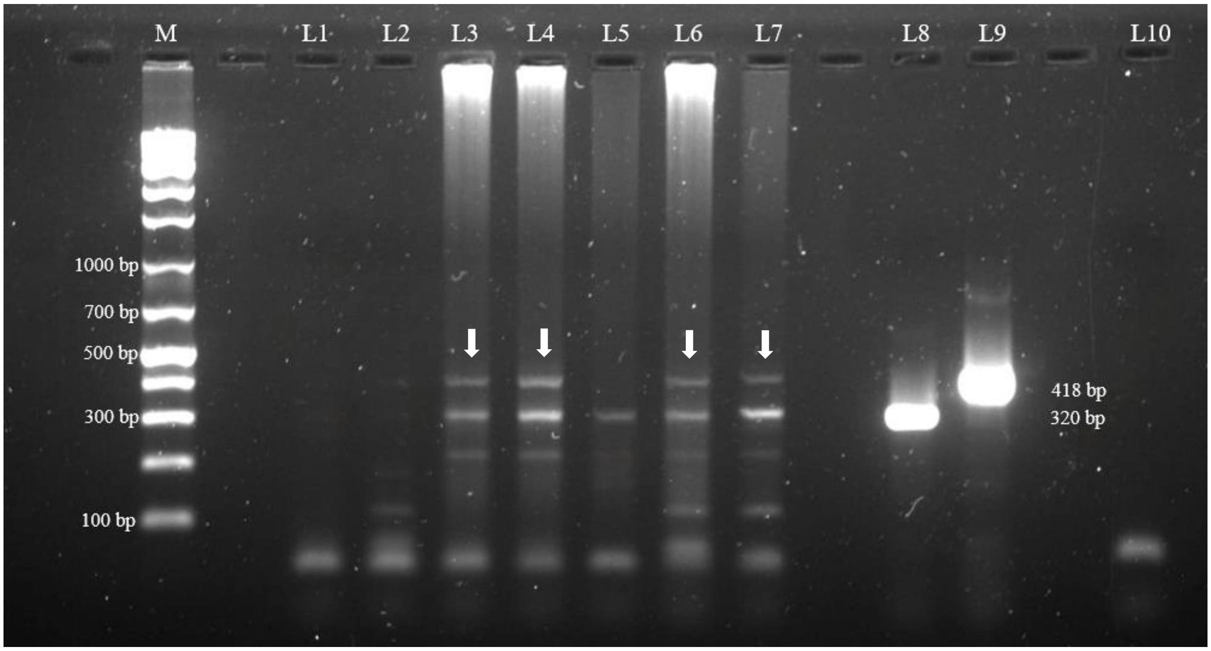

**Figure S3: Representative image of ITS1 PCR analysis from the DNA samples after ethanol precipitation of DNA.**

M: Marker; L1-L2: represents DNA from spleen cultures of mice infected with only LS after 2 m.p.i. followed by 1-month and 2-months cultures, respectively. L3: represents the same co-culture 10:1 infected spleen DNA sample, which was repeated with maximum template (not relevant). L4-L5: represents positive control DNA from LD and LS, respectively; L6: represents negative control (where L represents lane number), L1-L2 white arrow indicates the presence and detection of LS positive band in 1- and 2-months spleen culture at 2 m.p.i. for a LS infected mouse respectively.

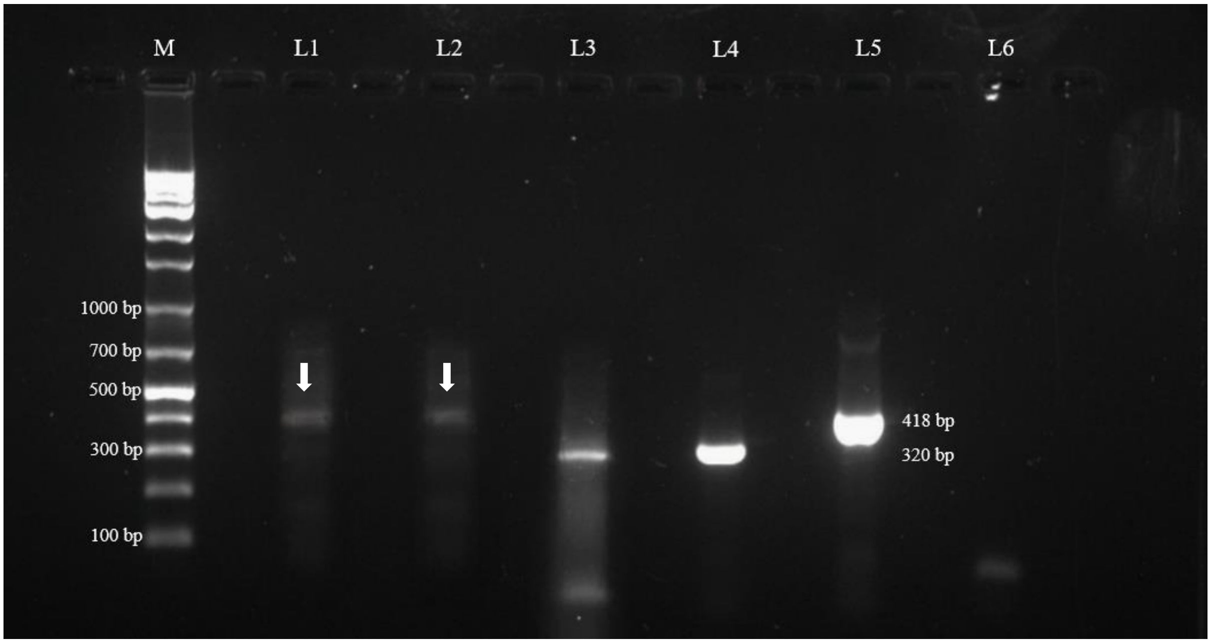

**Table S3: Estimated parasite copy number and (LD: LS) ratio in parasites recovered from infected bBALB/c mice following visceral organ culture.**

Spleen cultures established from BALB/c mice infected with LS, LD: LS co-infections (5:1 or 10:1), or AG83 laboratory isolate were analysed for parasite burden after the indicated duration of *in vivo* infection and subsequent *ex vivo* culture. Estimated parasite copy numbers were determined by ITS1 PCR followed by ImageJ analysis and are presented as approximate ranges.

| SAMPLE | INFECTION/CULTURE CONDITION | ESTIMATED COPY NUMBER | RATIO (LD: LS) |
|---|---|---|---|
| **LS** | 2 m.p.i. / 1-month culture | $10^4$-$10^5$ | |
| **LS** | 2 m.p.i./ 2 months culture | $10^4$ | |
| **5:1 Co-inf-LD** | A1-5 m.p.i./ 1-month culture | $10^5$-$10^6$ | 10:1 |
| **5:1 Co-inf-LS** | | $10^5$ | |
| **5:1 Co-inf-LD** | A2-5 m.p.i./ 1-month culture | $10^6$ | 10:1-1:1 |
| **5:1 Co-inf-LS** | | $10^5$- $10^6$ | |
| **10:1 Co-inf-LD** | 5 m.p.i./ 1-month culture | $10^4$ - $10^5$ | Only LD detected |
| **AG83-LD** | A1-5.5 m.p.i./ 1-month culture | $10^5$- $10^6$ | 1:1 |
| **AG83-LS** | | $10^5$- $10^6$ | |
| **AG83-LD** | A2-5.5 m.p.i. /1-month culture | $10^5$- $10^6$ | 10:1 |
| **AG83-LS** | | $10^5$ | |
| **8-LD-PC** | | .> $10^7$ | |
| **9-LS-PC** | | .> $10^7$ | |

**Figure S4: Representative image of LS DNA PCR and RT-PCR analysis from spleen culture samples of BALB/c mice infected with only LS at 2 m.p.i. followed by a 1-month culture.**

M: Marker; L1: represents spleen DNA; L2: represents positive control DNA from LS*;* L3: represents negative control. L4-L6: first round RT-PCR products from spleen RNA, positive control for Lepsey NLV1 and negative control respectively; L7-L9: represents second round PCR products from spleen RNA, positive control for Lepsey NLV1 and negative control respectively (where L represents lane number); L7 white arrow indicates the presence and detection of virus positive band in 1-month spleen culture at 2 m.p.i. for a LS infected mouse.

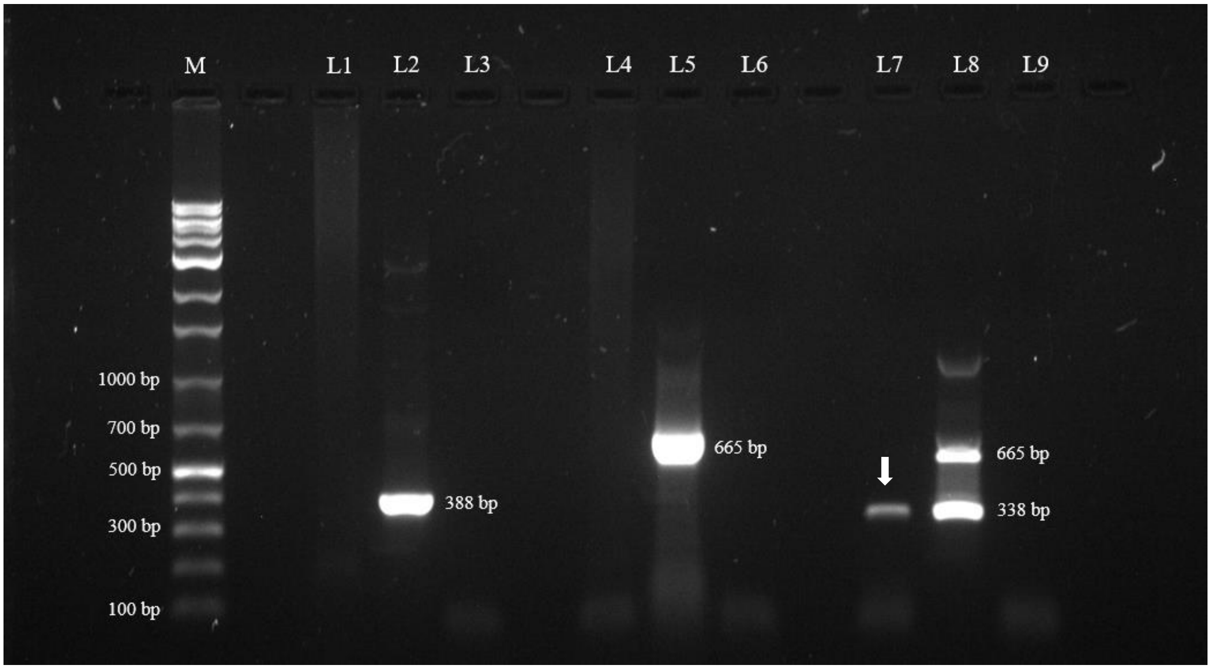

**Figure S5: Representative image of RT-PCR analysis for the detection of Lepsey NLV1 in splenocyte samples from infected BALB/c mice after 3 m.p.i.**

M: marker; L1-L4: represents first round PCR products from LD, LS, (LD: LS) 5:1 and 10:1 co-infected sample, respectively; L5 and L6: represents first round PCR products of positive control and negative control respectively. L7-L10: represents the second round PCR products from LD, LS, 5:1 and 10:1 co-infected sample, respectively; L11-L12: Second round PCR samples for positive control and negative control respectively (where L represents Lane number**);** L8-L10 white arrow indicates the presence and detection of virus positive band at 3 m.p.i. from mice infected with LS, 5:1 and 10:1 co-infection respectively.

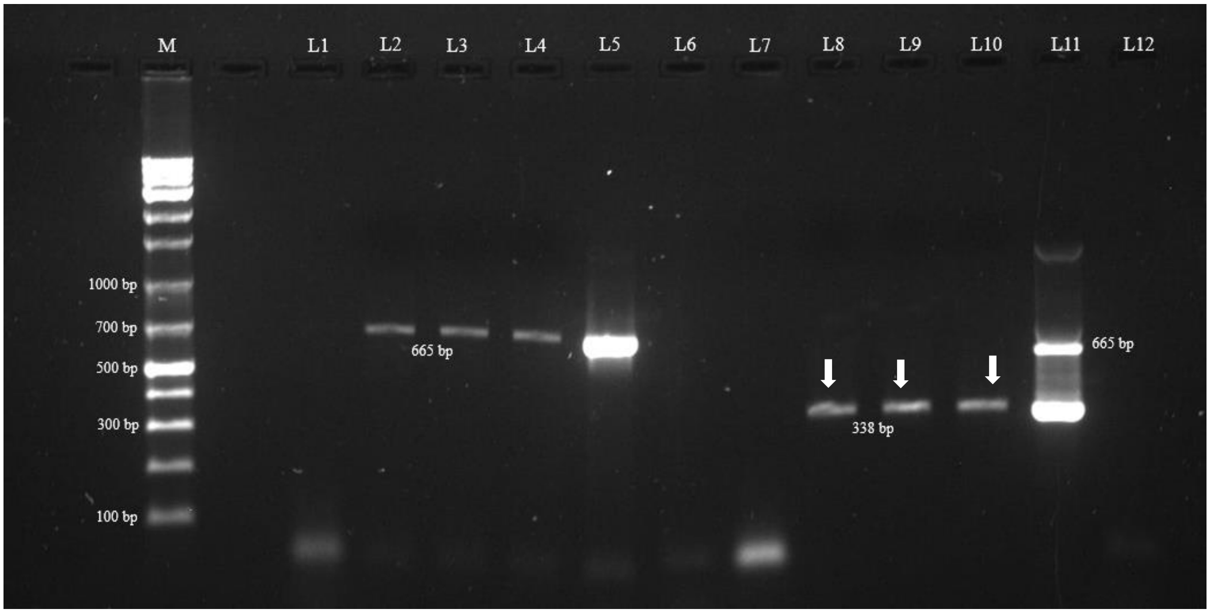

**Figure S6: Representative image of RT-PCR analysis of RNA from spleen samples of BALB/c mice infected with parasites**.

M: marker; L1: represents negative control for first round PCR; L2-L4: represents first round PCR products from spleen of mice infected with only LD after 1-m.p.i. followed by 13 days of tissue culture, (LD: LS) 10:1 co-infected spleen sample after 1-m.p.i. followed by 13 days of tissue culture; 2:1 co- infected spleen sample after 1-m.p.i. followed by 5 days of tissue culture respectively; L5: represents first round positive control from RNA from Lepsey NLV1; L6: second round PCR for negative control; L7-L9: represents second round PCR from same spleen of mice infected with only LD after 1-m.p.i. followed by 13 days of tissue culture, 10:1 co- infected spleen sample after 1-m.p.i. followed by 13 days of tissue culture; 2:1 co- infected spleen sample after 1-m.p.i. followed by 5 days of tissue culture respectively; L10: Second round PCR samples for positive control (where L represents Lane number**).** L8-L9 white arrow indicates the presence and detection of virus in 13 days and 5 days spleen tissue cultures at 1-m.p.i. for a 10:1 and 2:1 co- infected mouse.

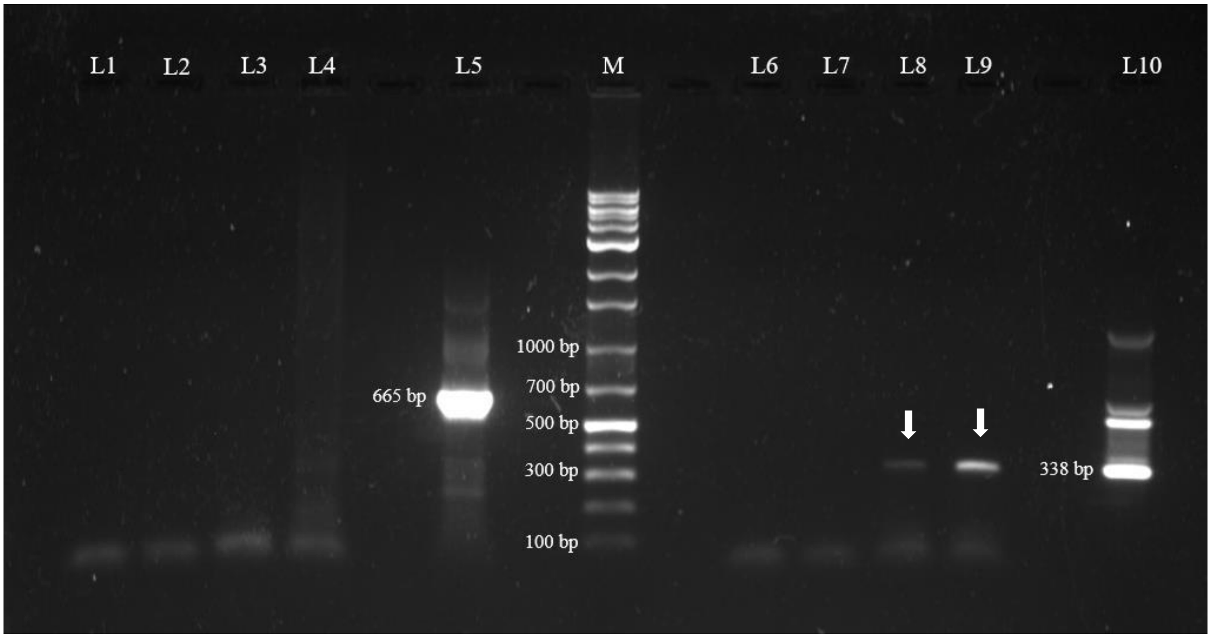

**Figure S7: Representative image of RT-PCR analysis for the detection of Lepsey NLV1 from BALB/c mice spleen samples infected with AG83 lab isolate**.

M: marker; L1-L2: represents first round PCR products from spleen tissue stored in RNA Later after 5.5 m.p.i. (representative of two different biological replicates); L3-L4: represents first round PCR products from the same samples after 5.5 m.p.i. followed by a 1-month tissue culture (representative of two different biological replicates); L5-L6: represents the first round PCR products of the positive control and negative control, respectively. L7-L8: represents the second round PCR of the same spleen samples directly from tissue (representative of two different biological replicates); L9-L10: represents the second round PCR of the same spleen samples after 1 month of tissue culture (representative of two different biological replicates); L11-L12: Second round PCR samples for positive control and negative control respectively (where L represents Lane number**).** White arrow at L9-L10 indicates the detection and presence of virus in 1-month spleen tissue cultures at 5.5 m.p.i. for a virus-positive AG83 lab isolate infected mice.

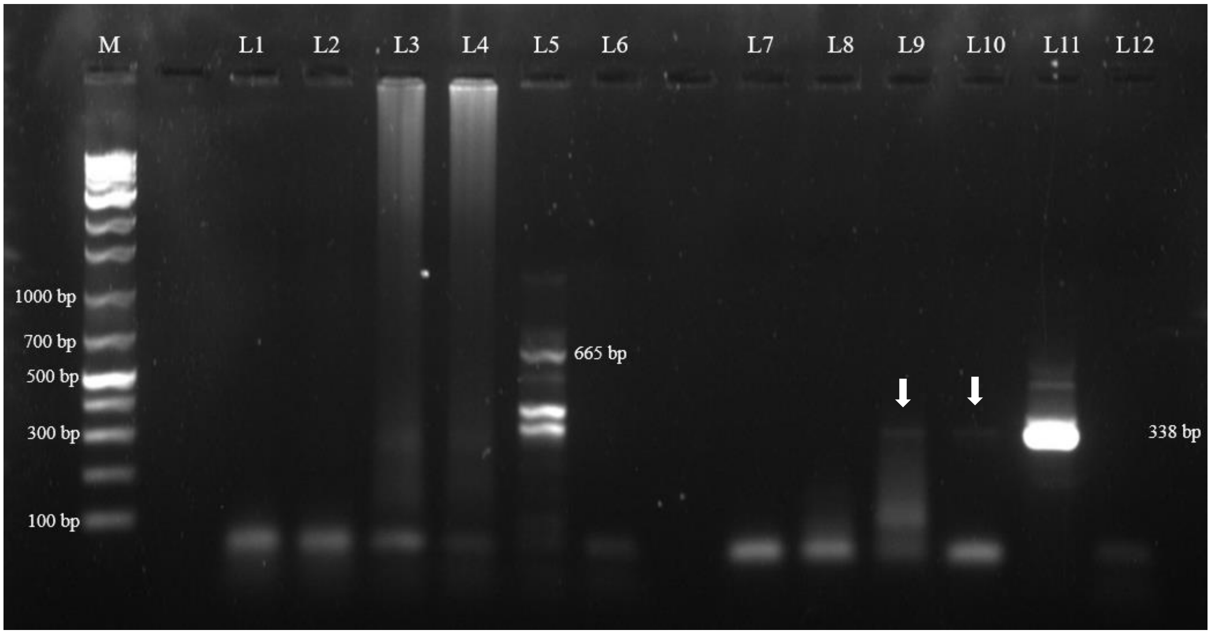

**Figure S8: Representative image of RT-PCR analysis from blood samples of BALB/c mice infected with parasites.**

M: marker; L1-L4: represents first PCR products for negative control, RNA samples of spleen infected with LD, 10:1 and 2:1 co-infection of LD: LS respectively after 1-m.p.i. L5-L8: represents the second round PCR products of the RNA samples of spleen infected with LD, 10:1 and 2:1 co-infection of LD: LS after 1-m.p.i. and negative control respectively, (where L represents Lane number**).** L7 white arrow suggests the presence and detection of virus in blood at 1-m.p.i. for a 2:1 co-infected mouse.

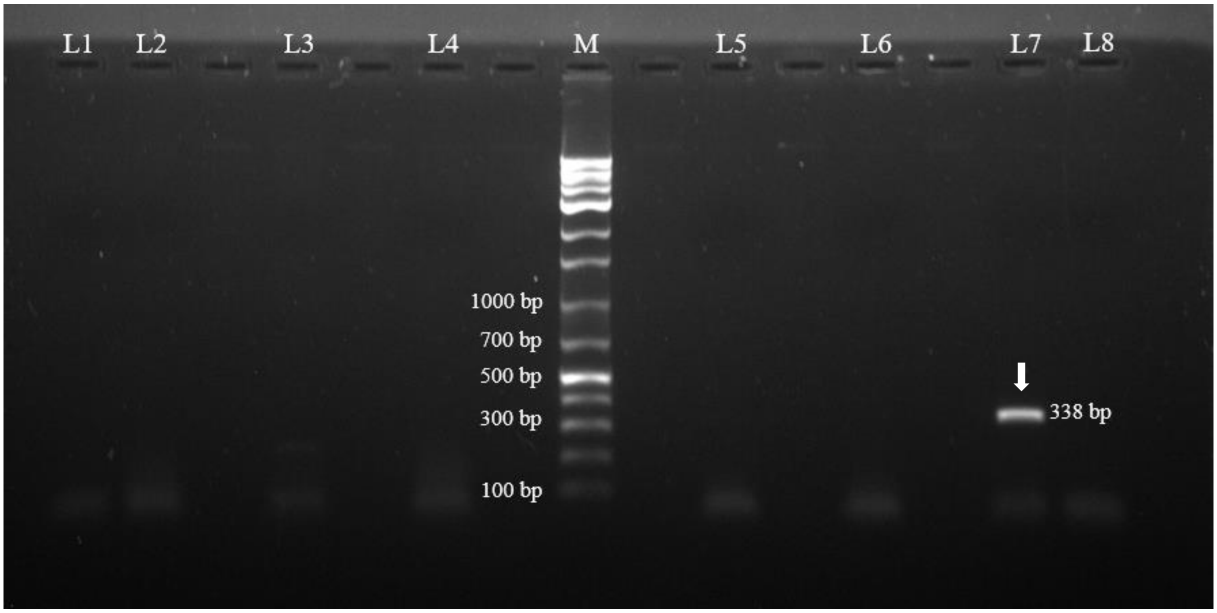

**Figure S9: Representative image of RT-PCR analysis from blood samples of BALB/c mice infected with parasites**.

M: marker; L1 and L3: represents first round PCR products from blood infected with LS and (LD: LS) 10:1 co-infection after 2 m.p.i. respectively. L2 and L4: represents the first round PCR products of same samples after 5 m.p.i. respectively. L5 and L6: represents positive and negative control respectively. L7 and L9: represents second round PCR products from blood infected with LS and 10:1 co-infection after 2 m.p.i. respectively. L8 and L10: represents the second round PCR products of same samples after 5 m.p.i. respectively; L11: Positive control for Lepsey NLV1; L12: Negative control (where L represents Lane number). L10 white arrow indicates the presence and detection of virus in blood at 5-m.p.i. for a 10:1 co- infected mouse.

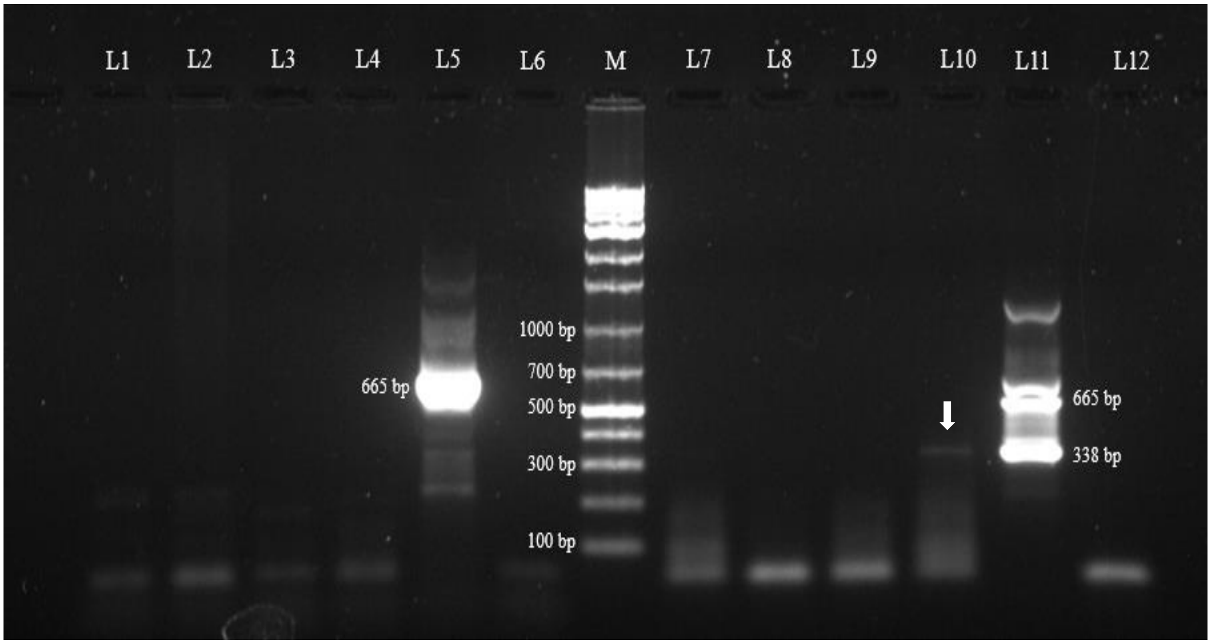

**Table S4. Multiple sequence alignment of the amplified Lepsey NLV1 fragment recovered from experimentally infected BALB/c mice with closely related Lepsey NLV1 sequences identified by BLAST analysis.**

The amplified Lepsey NLV1 sequences obtained from infected mouse samples were aligned with representative closely related Lepsey NLV1 sequences retrieved through BLAST analysis to assess sequence similarity. Sample 2L represents the spleen culture isolate obtained from a mouse infected with a 2:1 co-infection (LD: LS) and collected at 1 m.p.i. Sample L10 represents the spleen culture isolate from a mouse infected with a 10:1 co-infection (LD: LS) at 1 m.p.i. Sample 4TA1 represents a blood sample collected from a mouse infected with a 10:1 co-infection (LD: LS) at 5 m.p.i. Conserved nucleotides are indicated by alignment with homologous Lepsey NLV1 sequences, demonstrating high sequence identity between the experimentally recovered viral isolates and previously reported Lepsey NLV1 sequences.

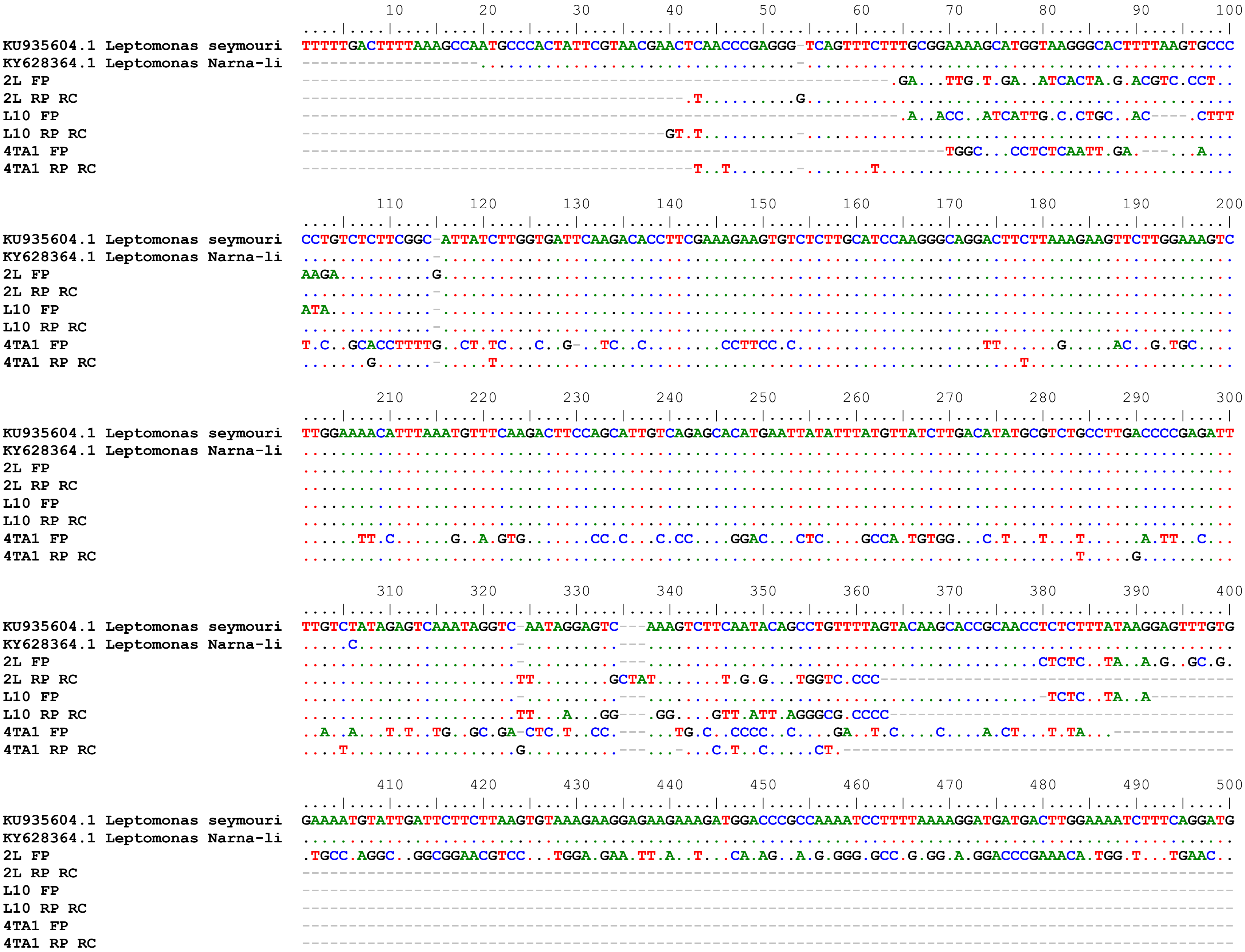
KU935604.1 Leptomonas seymouri
KY628364.1 Leptomonas Narna-li
2L FP
2L RP RC
L10 FP
L10 RP RC
4TA1 FP
4TA1 RP RC

```
                                       510       520       530       540       550       560       570       580       590       600
                                ....|....|....|....|....|....|....|....|....|....|....|....|....|....|....|....|....|....|....|....|
KU935604.1 Leptomonas seymouri  CAATTGCATCCGTGCAACTCTCCAAGGGGAAATCCCTTGAGAGGGTTCTCTCCGTTCTTACTACCCGGAACTTTCCAGAACCGGATGGTAAGTCTATACA
KY628364.1 Leptomonas Narna-li  ............................A.......................................................................
2L FP                            GT..CAGC.TGTG.GCTT..GTGGG....G.GGGAGGGACC.CCCCC.C.C..TGGGCGTA.TTTTATG.GACCGATAC.GGA.GG..GGT.GAGGG.GG
2L RP RC                        ----------------------------------------------------------------------------------------------------
L10 FP                          ----------------------------------------------------------------------------------------------------
L10 RP RC                       ----------------------------------------------------------------------------------------------------
4TA1 FP                         ----------------------------------------------------------------------------------------------------
4TA1 RP RC                      ----------------------------------------------------------------------------------------------------

                                       610       620       630       640       650       660       670       680       690       700
                                ....|....|....|....|....|....|....|....|....|....|....|....|....|....|....|....|....|....|....|....|
KU935604.1 Leptomonas seymouri  GAAGAAGAAAGTAGACTTCATAGGGATTATTTCTCAACAACCACCGAAAGGATGGAAGTCTGAGGAATATCGATCTCATTTCCTCGAATCTATAGAGGAG
KY628364.1 Leptomonas Narna-li  ....................................................................................................
2L FP                            .GG.T-----------------------------------------------------------------------------------------------
2L RP RC                        ----------------------------------------------------------------------------------------------------
L10 FP                          ----------------------------------------------------------------------------------------------------
L10 RP RC                       ----------------------------------------------------------------------------------------------------
4TA1 FP                         ----------------------------------------------------------------------------------------------------
4TA1 RP RC                      ----------------------------------------------------------------------------------------------------

                                       710       720       730       740       750       760       770       780       790       800
                                ....|....|....|....|....|....|....|....|....|....|....|....|....|....|....|....|....|....|....|....|
KU935604.1 Leptomonas seymouri  ATCGCAAATGAATGCGAATCCACTACAGTTTCCGATTGCCACATTTCTGTGACAGCCGCCGGCTCACTGAAGAAAACAGTGAGGGAGGGAGGAAAGTTCG
KY628364.1 Leptomonas Narna-li  ....................................................................................................
2L FP                            ----------------------------------------------------------------------------------------------------
2L RP RC                        ----------------------------------------------------------------------------------------------------
L10 FP                          ----------------------------------------------------------------------------------------------------
L10 RP RC                       ----------------------------------------------------------------------------------------------------
4TA1 FP                         ----------------------------------------------------------------------------------------------------
4TA1 RP RC                      ----------------------------------------------------------------------------------------------------

                                       810       820       830       840       850       860       870       880       890       900
                                ....|....|....|....|....|....|....|....|....|....|....|....|....|....|....|....|....|....|....|....|
KU935604.1 Leptomonas seymouri  CAGAGATGATCGAGGAAGTTAAAGGTTTCCTTGCGGAGACCCCTTCCAACGAGGAGCTCTACGAATTTGCAAATCTAAAATGGACTTGCAATACCAATGA
KY628364.1 Leptomonas Narna-li  ....................................................................................................
2L FP                            ----------------------------------------------------------------------------------------------------
2L RP RC                        ----------------------------------------------------------------------------------------------------
L10 FP                          ----------------------------------------------------------------------------------------------------
L10 RP RC                       ----------------------------------------------------------------------------------------------------
4TA1 FP                         ----------------------------------------------------------------------------------------------------
4TA1 RP RC                      ----------------------------------------------------------------------------------------------------

                                       910       920       930       940       950       960       970       980       990      1000
                                ....|....|....|....|....|....|....|....|....|....|....|....|....|....|....|....|....|....|....|....|
KU935604.1 Leptomonas seymouri  ACCAAGGTGGAAGACCTTCGGTATTATTGGTGAGATCAATCCCATGTCCGCTATGACTCTCTTTACCGAAGGAGTGGACTTCTTAAACGAAGTTCCAAAC
KY628364.1 Leptomonas Narna-li  ....................................................................................................
2L FP                            ----------------------------------------------------------------------------------------------------
2L RP RC                        ----------------------------------------------------------------------------------------------------
L10 FP                          ----------------------------------------------------------------------------------------------------
L10 RP RC                       ----------------------------------------------------------------------------------------------------
4TA1 FP                         ----------------------------------------------------------------------------------------------------
4TA1 RP RC                      ----------------------------------------------------------------------------------------------------
```

```
                                     1010      1020      1030      1040      1050      1060      1070      1080      1090      1100
                             ....|....|....|....|....|....|....|....|....|....|....|....|....|....|....|....|....|....|....|....|
KU935604.1 Leptomonas seymouri  CAGTACTTGGGTCCAATTGACTTTTCTGGAGGGTCACTTCCCCCTGGAATTATTCCAGAGGAGGATCTGTTTATAACAGATCTAAGACCTTTCAGAATGG
KY628364.1 Leptomonas Narna-li  ....................................................................................................
2L FP                           ----------------------------------------------------------------------------------------------------
2L RP RC                        ----------------------------------------------------------------------------------------------------
L10 FP                          ----------------------------------------------------------------------------------------------------
L10 RP RC                       ----------------------------------------------------------------------------------------------------
4TA1 FP                         ----------------------------------------------------------------------------------------------------
4TA1 RP RC                      ----------------------------------------------------------------------------------------------------

                                     1110      1120      1130      1140      1150      1160      1170      1180      1190      1200
                             ....|....|....|....|....|....|....|....|....|....|....|....|....|....|....|....|....|....|....|....|
KU935604.1 Leptomonas seymouri  GTCTTGGAGCCCAATTCGGTAATCAGCTTTTACTTTACTGCTGCGTCTTTTATAAAAAAGATGAGCTCCCTATGATTAGGGCCTCCCCTGTGTTAGAGGG
KY628364.1 Leptomonas Narna-li  ............................C.......................................................................
2L FP                           ----------------------------------------------------------------------------------------------------
2L RP RC                        ----------------------------------------------------------------------------------------------------
L10 FP                          ----------------------------------------------------------------------------------------------------
L10 RP RC                       ----------------------------------------------------------------------------------------------------
4TA1 FP                         ----------------------------------------------------------------------------------------------------
4TA1 RP RC                      ----------------------------------------------------------------------------------------------------

                                     1210      1220      1230      1240      1250      1260      1270      1280      1290      1300
                             ....|....|....|....|....|....|....|....|....|....|....|....|....|....|....|....|....|....|....|....|
KU935604.1 Leptomonas seymouri  AGGAGATAAAGTAAGATGGATAACGATGGCCTCATGGAGAGACCTCGTTATTCAACAAGCTGCAGCAACGATCTTTCGATCGTTGATGGAGAGCCATAAG
KY628364.1 Leptomonas Narna-li  ....................................................................................................
2L FP                           ----------------------------------------------------------------------------------------------------
2L RP RC                        ----------------------------------------------------------------------------------------------------
L10 FP                          ----------------------------------------------------------------------------------------------------
L10 RP RC                       ----------------------------------------------------------------------------------------------------
4TA1 FP                         ----------------------------------------------------------------------------------------------------
4TA1 RP RC                      ----------------------------------------------------------------------------------------------------

                                     1310      1320      1330      1340      1350      1360      1370      1380      1390      1400
                             ....|....|....|....|....|....|....|....|....|....|....|....|....|....|....|....|....|....|....|....|
KU935604.1 Leptomonas seymouri  GAAATGAAACCTATTTTCTCGCGTGCAAATCTTGCTTGGGTTTACCTAAACAAGGCACGTGAGATAGGACCTAGTGACATCTGTTACGTTTCGGACTACA
KY628364.1 Leptomonas Narna-li  ....................................................................................................
2L FP                           ----------------------------------------------------------------------------------------------------
2L RP RC                        ----------------------------------------------------------------------------------------------------
L10 FP                          ----------------------------------------------------------------------------------------------------
L10 RP RC                       ----------------------------------------------------------------------------------------------------
4TA1 FP                         ----------------------------------------------------------------------------------------------------
4TA1 RP RC                      ----------------------------------------------------------------------------------------------------

                                     1410      1420      1430      1440      1450      1460      1470      1480      1490      1500
                             ....|....|....|....|....|....|....|....|....|....|....|....|....|....|....|....|....|....|....|....|
KU935604.1 Leptomonas seymouri  GTTCTGCAACGGACACAGTTGACAGGGAGTTTGCTGAGTTTATCTTAACAAACTTTATCAAGCGGTTCCGCAATAGACTTTCTGATCCTCTCCTCAATTT
KY628364.1 Leptomonas Narna-li  ....................................................................................................
2L FP                           ----------------------------------------------------------------------------------------------------
2L RP RC                        ----------------------------------------------------------------------------------------------------
L10 FP                          ----------------------------------------------------------------------------------------------------
L10 RP RC                       ----------------------------------------------------------------------------------------------------
4TA1 FP                         ----------------------------------------------------------------------------------------------------
4TA1 RP RC                      ----------------------------------------------------------------------------------------------------
```

```
                                         1510      1520      1530      1540      1550      1560      1570      1580      1590      1600
                                 ....|....|....|....|....|....|....|....|....|....|....|....|....|....|....|....|....|....|....|....|
KU935604.1 Leptomonas seymouri   CTTAGAATTAGGGATTAGGAATGCAGTAAGTCCCAAGGTTGTTATCTTTCCGACAGGAGAGAGAATAACCTCAAGCAGAGGCGTCTTCATGGGTGAACCG
KY628364.1 Leptomonas Narna-li   ....................................................................................................
2L FP                            ----------------------------------------------------------------------------------------------------
2L RP RC                         ----------------------------------------------------------------------------------------------------
L10 FP                           ----------------------------------------------------------------------------------------------------
L10 RP RC                        ----------------------------------------------------------------------------------------------------
4TA1 FP                          ----------------------------------------------------------------------------------------------------
4TA1 RP RC                       ----------------------------------------------------------------------------------------------------

                                         1610      1620      1630      1640      1650      1660      1670      1680      1690      1700
                                 ....|....|....|....|....|....|....|....|....|....|....|....|....|....|....|....|....|....|....|....|
KU935604.1 Leptomonas seymouri   ATGAGTAAAGTTATTCTTACACTCATCATGTTCACGATAGGAAAGGCCGCAAAATCAATATACAAATTGAGATTCCCAAAGTCAATGGAGAAATTGACAT
KY628364.1 Leptomonas Narna-li   ....................................................................................................
2L FP                            ----------------------------------------------------------------------------------------------------
2L RP RC                         ----------------------------------------------------------------------------------------------------
L10 FP                           ----------------------------------------------------------------------------------------------------
L10 RP RC                        ----------------------------------------------------------------------------------------------------
4TA1 FP                          ----------------------------------------------------------------------------------------------------
4TA1 RP RC                       ----------------------------------------------------------------------------------------------------

                                         1710      1720      1730      1740      1750      1760      1770      1780      1790      1800
                                 ....|....|....|....|....|....|....|....|....|....|....|....|....|....|....|....|....|....|....|....|
KU935604.1 Leptomonas seymouri   TTTGGGCACCAGGTGATGACCTGGTGGCGACTGGCCCTACCGAATATATTGACATATATTCCGAACTCGCCAAGATCCTTGGTCAGATTCTTAACCACTC
KY628364.1 Leptomonas Narna-li   ....................................................................................................
2L FP                            ----------------------------------------------------------------------------------------------------
2L RP RC                         ----------------------------------------------------------------------------------------------------
L10 FP                           ----------------------------------------------------------------------------------------------------
L10 RP RC                        ----------------------------------------------------------------------------------------------------
4TA1 FP                          ----------------------------------------------------------------------------------------------------
4TA1 RP RC                       ----------------------------------------------------------------------------------------------------

                                         1810      1820      1830      1840      1850      1860      1870      1880      1890      1900
                                 ....|....|....|....|....|....|....|....|....|....|....|....|....|....|....|....|....|....|....|....|
KU935604.1 Leptomonas seymouri   AAAGGTGTTTAAGTCACGCACAGTTTTTAAACTGTGTGAGCAATGGTTTTGGGTGCCAGGGCTGAAAAGCTCTGTTGGCACCTGGGCCATCACTACTGAT
KY628364.1 Leptomonas Narna-li   ....................................................................................................
2L FP                            ----------------------------------------------------------------------------------------------------
2L RP RC                         ----------------------------------------------------------------------------------------------------
L10 FP                           ----------------------------------------------------------------------------------------------------
L10 RP RC                        ----------------------------------------------------------------------------------------------------
4TA1 FP                          ----------------------------------------------------------------------------------------------------
4TA1 RP RC                       ----------------------------------------------------------------------------------------------------

                                         1910      1920      1930      1940      1950      1960      1970      1980      1990      2000
                                 ....|....|....|....|....|....|....|....|....|....|....|....|....|....|....|....|....|....|....|....|
KU935604.1 Leptomonas seymouri   CCAGGGAAGTATAGGGAATCCGCATGGGTAGACACCGTGAAATTAAAACTTCTCGGTGCCATGAGTCTGAGTAATCATAGTCACTTTGAAGAGAGAAACG
KY628364.1 Leptomonas Narna-li   ....................................................................................................
2L FP                            ----------------------------------------------------------------------------------------------------
2L RP RC                         ----------------------------------------------------------------------------------------------------
L10 FP                           ----------------------------------------------------------------------------------------------------
L10 RP RC                        ----------------------------------------------------------------------------------------------------
4TA1 FP                          ----------------------------------------------------------------------------------------------------
4TA1 RP RC                       ----------------------------------------------------------------------------------------------------
```

```
                                         2010      2020      2030      2040      2050      2060      2070      2080      2090      2100
                                ....|....|....|....|....|....|....|....|....|....|....|....|....|....|....|....|....|....|....|....|
KU935604.1 Leptomonas seymouri  AGGTTATTGGGAAGGCGAAAGCCTTATCCAAAATTCTTCGTTGGCTCCCTAGTGACACTTACCCTTTAGAGTATAAAAAGCTCTTAAGGACTTGGTTTTT
KY628364.1 Leptomonas Narna-li  ....................................................................................................
2L FP                           ----------------------------------------------------------------------------------------------------
2L RP RC                        ----------------------------------------------------------------------------------------------------
L10 FP                          ----------------------------------------------------------------------------------------------------
L10 RP RC                       ----------------------------------------------------------------------------------------------------
4TA1 FP                         ----------------------------------------------------------------------------------------------------
4TA1 RP RC                      ----------------------------------------------------------------------------------------------------

                                         2110      2120      2130      2140      2150      2160      2170      2180      2190      2200
                                ....|....|....|....|....|....|....|....|....|....|....|....|....|....|....|....|....|....|....|....|
KU935604.1 Leptomonas seymouri  GTGTAGGTTTGAACCTAGATTACCAAGAACAGACTCAAAGACCTTTGCTTTCCTTATGTTACCCGGACACCTAGGAGGTTTTGACCTCCTGCTAGATGAC
KY628364.1 Leptomonas Narna-li  .................A..................................................................................
2L FP                           ----------------------------------------------------------------------------------------------------
2L RP RC                        ----------------------------------------------------------------------------------------------------
L10 FP                          ----------------------------------------------------------------------------------------------------
L10 RP RC                       ----------------------------------------------------------------------------------------------------
4TA1 FP                         ----------------------------------------------------------------------------------------------------
4TA1 RP RC                      ----------------------------------------------------------------------------------------------------

                                         2210      2220      2230      2240      2250      2260      2270      2280      2290      2300
                                ....|....|....|....|....|....|....|....|....|....|....|....|....|....|....|....|....|....|....|....|
KU935604.1 Leptomonas seymouri  CAGGAGATAAGAGAGGCATACCAAAAAGTGAGTTCCTTCTCTAGAATCTTATTTAACAACTACTATGATGATTTCATAGTTGGAAGTTATCTAAGAAACA
KY628364.1 Leptomonas Narna-li  ....................................................................................................
2L FP                           ----------------------------------------------------------------------------------------------------
2L RP RC                        ----------------------------------------------------------------------------------------------------
L10 FP                          ----------------------------------------------------------------------------------------------------
L10 RP RC                       ----------------------------------------------------------------------------------------------------
4TA1 FP                         ----------------------------------------------------------------------------------------------------
4TA1 RP RC                      ----------------------------------------------------------------------------------------------------

                                         2310      2320      2330      2340      2350      2360      2370      2380      2390      2400
                                ....|....|....|....|....|....|....|....|....|....|....|....|....|....|....|....|....|....|....|....|
KU935604.1 Leptomonas seymouri  TGTTGAGAAATAGGTCCTATAGGGGTTTTGCCCTATCGGAAGACTATAATCTCACCGTAGAGAAGCTGGTCAAGTATTATACTGACCTGCTTGAGGAAGC
KY628364.1 Leptomonas Narna-li  ............................C.......................................................................
2L FP                           ----------------------------------------------------------------------------------------------------
2L RP RC                        ----------------------------------------------------------------------------------------------------
L10 FP                          ----------------------------------------------------------------------------------------------------
L10 RP RC                       ----------------------------------------------------------------------------------------------------
4TA1 FP                         ----------------------------------------------------------------------------------------------------
4TA1 RP RC                      ----------------------------------------------------------------------------------------------------

                                         2410      2420      2430      2440      2450      2460      2470      2480      2490      2500
                                ....|....|....|....|....|....|....|....|....|....|....|....|....|....|....|....|....|....|....|....|
KU935604.1 Leptomonas seymouri  ACCTTATGAGGAGTACAAAGACTCCAAGGTGGAGGGTCTCTCATTTAGAGAGACCGACCGTGCCCTTTTTAAGGCAGGCTTCCTCACGGAAAACAGGATA
KY628364.1 Leptomonas Narna-li  ....................................................................................................
2L FP                           ----------------------------------------------------------------------------------------------------
2L RP RC                        ----------------------------------------------------------------------------------------------------
L10 FP                          ----------------------------------------------------------------------------------------------------
L10 RP RC                       ----------------------------------------------------------------------------------------------------
4TA1 FP                         ----------------------------------------------------------------------------------------------------
4TA1 RP RC                      ----------------------------------------------------------------------------------------------------
```

```
                                     2510      2520      2530      2540      2550      2560      2570      2580      2590      2600
                             ....|....|....|....|....|....|....|....|....|....|....|....|....|....|....|....|....|....|....|....|
KU935604.1 Leptomonas seymouri CGTGATATAGTATCACGTCCTCTCGCTTTTGGAAGCATATGGGGTAGAACGTTGAAACATTCACCCTATAATACCACTCCTATCTCTGTACGTCTCGCCA
KY628364.1 Leptomonas Narna-li ....................................................................................................
2L FP                         ----------------------------------------------------------------------------------------------------
2L RP RC                      ----------------------------------------------------------------------------------------------------
L10 FP                        ----------------------------------------------------------------------------------------------------
L10 RP RC                     ----------------------------------------------------------------------------------------------------
4TA1 FP                       ----------------------------------------------------------------------------------------------------
4TA1 RP RC                    ----------------------------------------------------------------------------------------------------

                                     2610      2620      2630      2640      2650      2660      2670      2680      2690      2700
                             ....|....|....|....|....|....|....|....|....|....|....|....|....|....|....|....|....|....|....|....|
KU935604.1 Leptomonas seymouri AACTCTGGAAGAATCTGACGGGAGATACAGAACGTTGGAAGAATATTTCATTGAAAGGTGAAATATCCGAAGACGATTTTGTAGTCCTGTGTAAAGAGAA
KY628364.1 Leptomonas Narna-li ....................................................................................................
2L FP                         ----------------------------------------------------------------------------------------------------
2L RP RC                      ----------------------------------------------------------------------------------------------------
L10 FP                        ----------------------------------------------------------------------------------------------------
L10 RP RC                     ----------------------------------------------------------------------------------------------------
4TA1 FP                       ----------------------------------------------------------------------------------------------------
4TA1 RP RC                    ----------------------------------------------------------------------------------------------------

                                     2710      2720      2730      2740      2750      2760      2770      2780      2790      2800
                             ....|....|....|....|....|....|....|....|....|....|....|....|....|....|....|....|....|....|....|....|
KU935604.1 Leptomonas seymouri TAAAGGTCCAGTACTCAAGGTCTACCGTCGAAACGATAGTCTTGAAGATGAAGTCCTCATAGGGCTTCCTTCTATGAAGGTAAGCCTCGGCATATTGCCC
KY628364.1 Leptomonas Narna-li ....................................................................................................
2L FP                         ----------------------------------------------------------------------------------------------------
2L RP RC                      ----------------------------------------------------------------------------------------------------
L10 FP                        ----------------------------------------------------------------------------------------------------
L10 RP RC                     ----------------------------------------------------------------------------------------------------
4TA1 FP                       ----------------------------------------------------------------------------------------------------
4TA1 RP RC                    ----------------------------------------------------------------------------------------------------

                                     2810      2820      2830      2840      2850      2860      2870      2880      2890      2900
                             ....|....|....|....|....|....|....|....|....|....|....|....|....|....|....|....|....|....|....|....|
KU935604.1 Leptomonas seymouri CATGATGGATACATCTCTTGGTAGTGGATCACTCTTTAGGCTCTTTACCAGTTGTATTTCTAAATACTTCTGGTTTCTGTGTGGGCTTTCAGTTTAAACT
KY628364.1 Leptomonas Narna-li ....................................................C.......................................--------
2L FP                         ----------------------------------------------------------------------------------------------------
2L RP RC                      ----------------------------------------------------------------------------------------------------
L10 FP                        ----------------------------------------------------------------------------------------------------
L10 RP RC                     ----------------------------------------------------------------------------------------------------
4TA1 FP                       ----------------------------------------------------------------------------------------------------
4TA1 RP RC                    ----------------------------------------------------------------------------------------------------

                                     2910      2920
                             ....|....|....|....|
KU935604.1 Leptomonas seymouri GAAAGCCCACACAGAAACCA
KY628364.1 Leptomonas Narna-li --------------------
2L FP                         --------------------
2L RP RC                      --------------------
L10 FP                        --------------------
L10 RP RC                     --------------------
4TA1 FP                       --------------------
4TA1 RP RC                    --------------------
```

**Figure S10: Representative microscopic views (1000X) of Giemsa-stained histological stamp smear of Spleen and Liver from mice at 7-m.p.i.**

Histological stamp smears of visceral organs **(A)** Spleen and **(B)** Liver from BALB/c mice infected with LD, LS, virus-positive AG83 and 10:1 co-infection of (LD: LS). Control mice were injected with sterile 1X PBS.

**(A)**

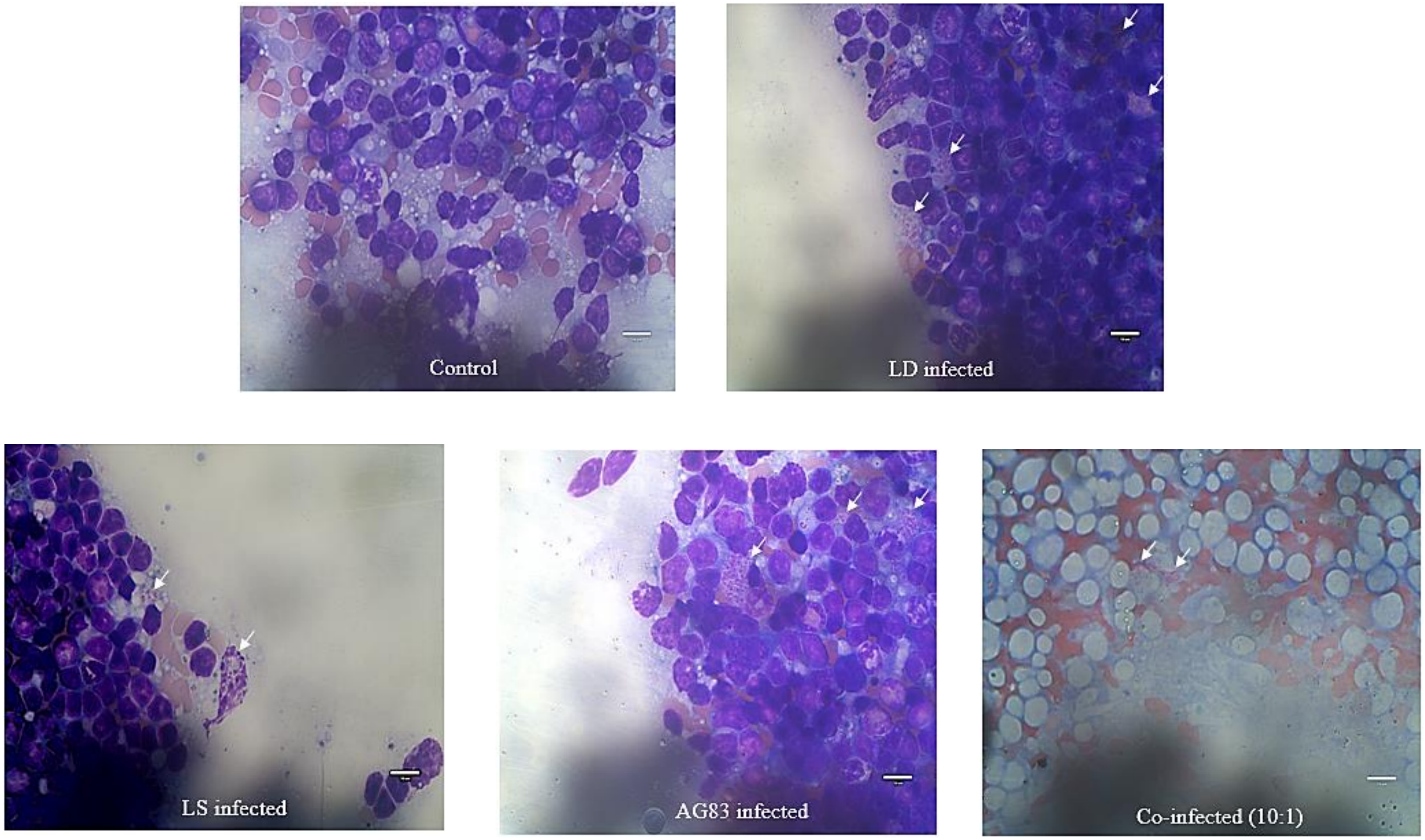

(B)

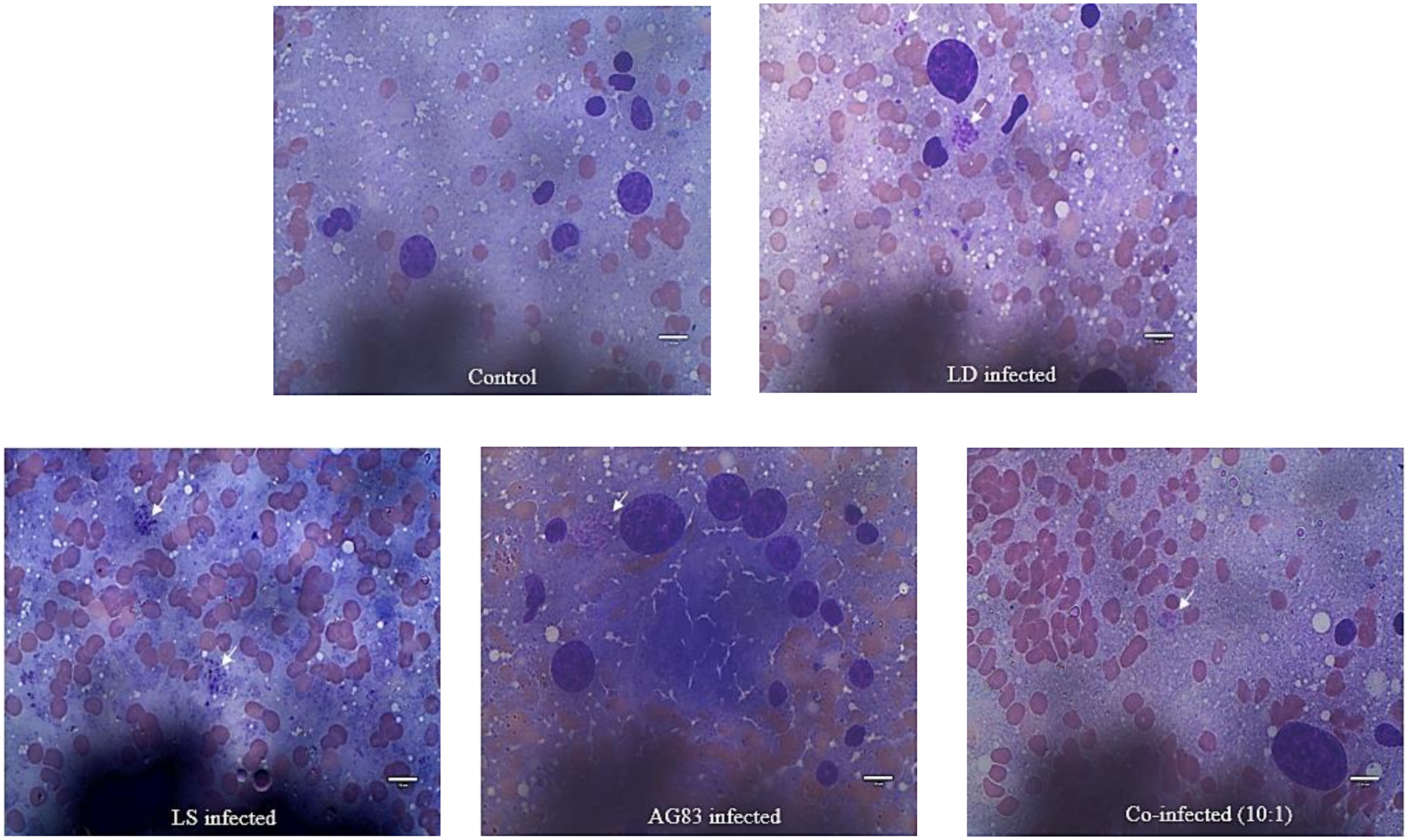
Control
LD infected
LS infected
AG83 infected
Co-infected (10:1)
